# A membrane homeostatic response to lipid overload coordinates fatty acid metabolism

**DOI:** 10.64898/2026.06.10.731409

**Authors:** Xiaojun Xiang, Yohannes A. Ambaw, Ozlem Tok, Sheng Hui, Wei-chun Tang, Arda Mizrak, Qingyue Zhang, Luke Cohen-Abeles, Robert V. Farese, Tobias C. Walther

**Author notes:** **Contact information:** Corresponding authors: Tobias C. Walther and Robert V. Farese Jr. These authors contributed equally (Robert V. Farese, Tobias C. Walther).

## Abstract

Excess fatty acids can disrupt membrane and organelle function. Cells buffer fatty acid toxicity by synthesizing and storing triglycerides (TGs) in lipid droplets, but their capacity for TG storage is limited. Here, we impaired TG synthesis in hepatocytes and identified adaptive pathways that restore homeostasis during lipid overload. Transcription responds by activating fatty acid oxidation through peroxisome proliferator-activated receptors, and by suppressing synthesis of new, unsaturated fatty acids through sterol regulatory element-binding protein 1 (SREBP1). Mechanistically, SREBP1 appears to respond to changes in ER membrane fluidity. These findings reveal a homeostatic system monitoring and maintaining the ER membrane, coordinating fatty acid synthesis and oxidation, a finding with broad implications for understanding lipid physiology and developing TG synthesis inhibitors for treating fatty liver disease.

## Main

Fatty acids are essential for cellular function. They serve as components of membrane phospholipids, precursors of signaling molecules, and reservoirs of metabolic energy. However, excessive fatty acid accumulation is toxic, impairing organelle and cell function by disrupting signaling pathways and membrane properties^1–3^. Such lipotoxicity is thought to contribute to metabolic diseases, such as metabolic dysfunction-associated steatotic liver disease^4,5^.

Cells buffer excess fatty acids by diverting them into triglycerides (TGs), which are relatively inert and can be packaged into lipid droplets (LDs), away from membranes^6–8^. TG synthesis is catalyzed by the evolutionarily unrelated diacylglycerol acyltransferases (DGATs), DGAT1 and DGAT2^9,10^. These enzymes are partially redundant but have distinct properties. DGAT1 is an endoplasmic reticulum (ER) enzyme that preferentially esterifies exogenous fatty acids, whereas DGAT2 localizes to both the ER and LDs and is more closely coupled to *de novo* lipogenesis^11–13^. Despite their protective roles in esterifying fatty acids, the cellular capacity for TG synthesis and storage is limited.

We therefore hypothesized that cells activate homeostatic mechanisms to restore fatty acid balance to prevent lipotoxicity when TG synthesis capacity is exceeded. Several observations suggest such a response exists. Fatty acids are ligands activating transcription factors, such as peroxisome proliferator-activated receptor α (PPARα), a major regulator of fatty acid oxidation^14,15^. Additionally, altering TG synthesis modulates transcriptional effects mediated by sterol regulatory element binding protein1 (SREBP1), a master regulator of fatty acid synthesis and saturation^16,17^ . In hepatocytes, increased TG synthesis via DGAT overexpression increases expression of SREBP1 and its lipogenic target genes^18^, whereas impaired DGAT activity suppresses SREBP1 activity^19–21^.

These findings suggest that SREBP1 may respond to changes in fatty acid flux, although the underlying mechanisms remain unclear. Polyunsaturated fatty acids (PUFAs) suppress SREBP1 activity^22^, possibly via mechanisms that alter liver X receptor (LXR)-mediated transcriptional activation of fatty acid synthesis^23^ or via other signaling pathways^24^. Additionally, inhibition of TG synthesis in hepatocytes was reported to alter ER phospholipid composition, including phosphatidylethanolamine (PE) accumulation, which was proposed to suppress fatty acids synthesis^25^. The physicochemical basis of SREBP1 regulation by all the different lipid metabolism changes has remained mysterious since the discovery of these transcription factors some 30 years ago^26^.

Here, we tested whether cells engage an integrated homeostatic response to maintain fatty acid balance when TG synthesis is impaired, analogous to stress-response systems, such as the integrated stress response (ISR)^27,28^ or the heat shock response^29^, that preserve other aspects of homeostasis. By chemically inhibiting DGAT activity in hepatocytes, we identified a coordinated feedback program that enhanced fatty acid oxidation while suppressing lipogenesis in response to limited TG synthesis capacity. Further, we dissected the underlying mechanisms for this response, providing new insights into ER membrane homeostasis.

## Results

### Impairing TG synthesis triggers similar transcriptional responses in fatty acid metabolism in murine liver and in human hepatocytes

The liver is the primary organ for handling fatty acids from multiple, different sources. Murine hepatocytes express both DGAT1 and DGAT2 enzymes for TG synthesis^30,31^. DGAT2 expression is directly correlated to expression of selected lipogenic genes^18–21^. To better understand how the liver responds to compromised TG synthesis, we performed RNA-seq analyses of gene expression in liver tissues from wild-type (WT) and liver-specific DGAT2 knockout (Liv-DGAT2KO) mice^21^. This analysis revealed an overall stable gene expression profile, and several strongly downregulated genes in Liv-DGAT2KO, including several regulated by SREBP1, such as *Scd1*, *Pltp*, and *Pnpla3* (Fig. 1a, b). Notably, the strongest downregulation observed was for the Δ-9 desaturase, *Scd1*.

**Figure 1.**
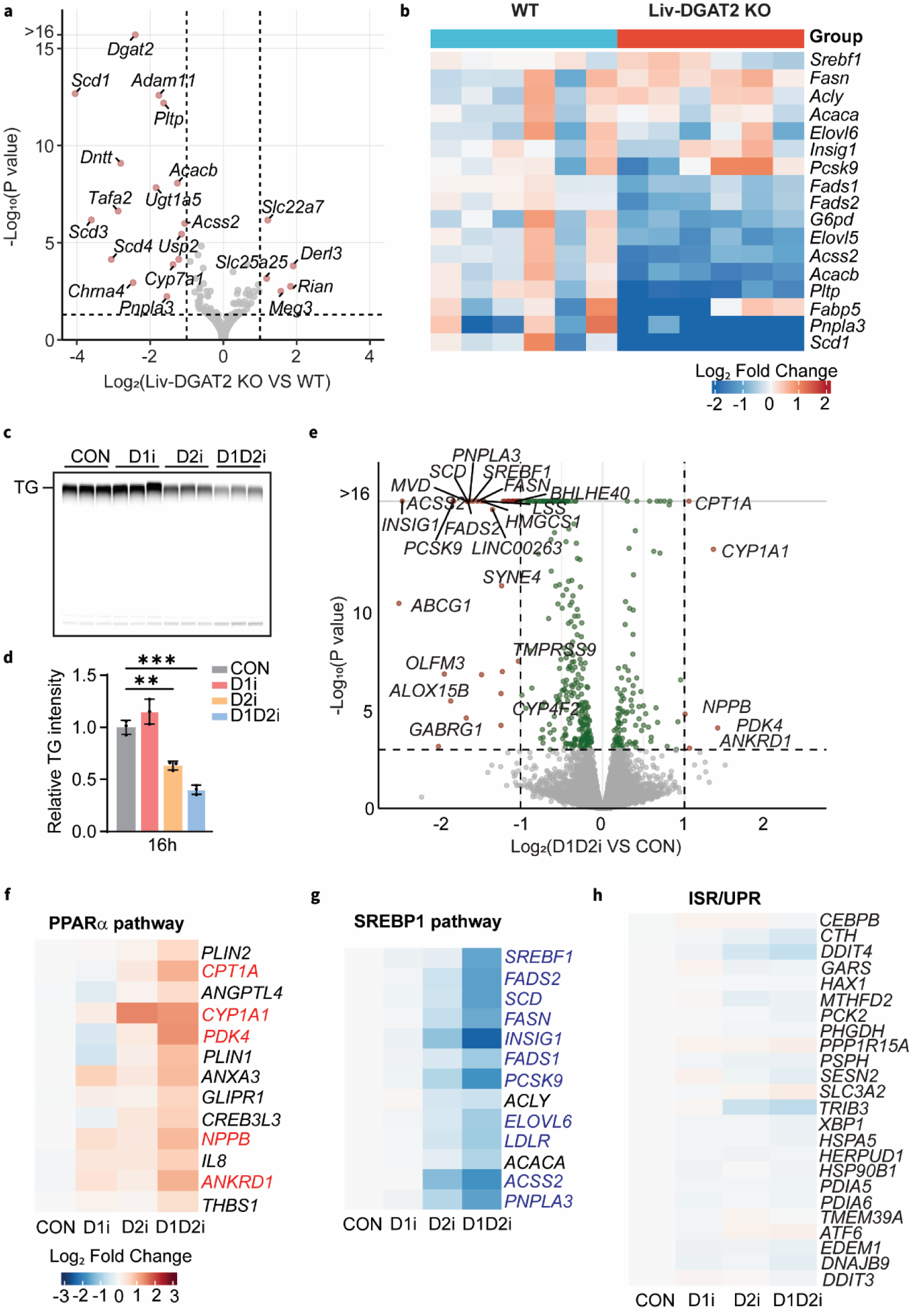
Inhibiting TG synthesis triggers a transcriptional response in murine liver and Huh-7 hepatoma cells. **(a)** Volcano plot showing differentially expressed genes in liver tissues between WT and liver-specific DGAT2 KO mice. Genes with adjusted p values < 0.05 and absolute fold-change > 1 were considered significantly differentially expressed. n=6 for each group. **(b)** Heatmap showing relative gene expression levels of genes involved in the SREBP1 pathway derived from RNA-seq analysis in liver tissues. **(c)** Incorporation of [^14^C] oleic acid into TG in Huh-7 cells treated with DMSO (CON), DGAT1i (D1i, 10 µM), DGAT2i (D2i, 10 µM) or both (D1D2i, 10 µM each) for 16 h was measured by extracting lipids from cells and separating them by the TLC. **(d)** Bar graph showing the quantification of labeled TG accumulation relative to control group. Error bars show mean ± SD of three independent experiments. Statistical significance was evaluated by unpaired two tailed *t*-test. ∗∗p < 0.01; ∗∗∗p < 0.001. **(e)** Volcano plot showing differentially expressed genes between combined DGAT1 and DGAT2 inhibition (D1D2i) and DMSO (CON) conditions. Genes with adjusted p values < 0.05 and absolute fold-change > 1 were considered significantly differentially expressed and shown in red. Genes with adjusted p values < 0.05 and absolute fold-change < 1 were shown in green. n = 3 for each treatment. **(f)** Heatmap showing relative gene expression levels of genes involved in the PPARα pathway derived from RNA-seq analysis. Genes significantly upregulated in D1D2i versus CON are shown in red. **(g)** Heatmap showing relative gene expression levels of genes involved in the SREBP1 pathway derived from RNA-seq analysis. Huh-7 cells were treated with DMSO (CON), DGAT1 inhibitor (D1i, 10 µM), DGAT2 inhibitor (D2i, 10 µM), or combined DGAT1 and DGAT2 inhibition (D1D2i, 10 µM each) for 16 h. Genes significantly downregulated in D1D2i versus CON are shown in blue. **(h)** Heatmap showing relative gene expression levels of integrated stress response (ISR) and unfolded protein response (UPR) marker genes after 16 h of DGAT inhibitors treatments.

To investigate the mechanism underlying this regulation in response to impaired TG synthesis, we established a cellular model for this response and selected the human hepatocellular carcinoma Huh-7 cell line for our analyses because it originates from liver and expresses both DGAT1 and DGAT2^32^. Inhibiting DGAT1 had no significant effect on TG synthesis over 16 h of incubation with C^14^-labeled oleate, whereas DGAT2 inhibition decreased the TG synthesis rate by ∼50%. When both enzymes were inhibited, the TG synthesis rate decreased to ∼30% of controls (Fig. 1c, d). Thus, in these cells, both enzymes can synthesize TG, but DGAT2 function cannot be fully compensated by DGAT1.

The transcriptional response of cells to inhibition of DGATs revealed two classes of genes that were either induced or repressed after DGAT1/2 inhibition (Fig. 1e). Among upregulated genes, those encoding enzymes required for efficient fatty acid catabolism stood out, including *CPT1A* and *PDK4*, encoding carnitine palmitoyltransferase and pyruvate dehydrogenase kinase, respectively. Genes downregulated in cells with DGAT2 or combined DGAT1/2 inhibition were enriched for fatty acid and cholesterol synthesis, including fatty acid synthase (*FASN*), stearoyl-CoA desaturase 1 (*SCD1*), and fatty acid desaturases (*FADS1* and *FADS2*) (Fig. 1e). This pattern was also found—and accentuated—when cells were incubated with oleic acid-containing medium, which enhances fatty acid flux into the TG synthesis pathway (Extended Data Fig. 1a).

*CPT1A* expression is upregulated by PPAR transcription factors^33^, whereas *FASN*, *SCD1*, and *FADS* genes are regulated by SREBP1^34,35^. We therefore more broadly examined the expression of PPARα and SREBP1 target genes in the RNA-seq data. Heatmaps of gene expression changes showed that DGAT2 and combined DGAT1/2 inhibition upregulated PPARα targets but downregulated SREBP1 targets (Fig. 1f, g and Extended Data Fig. 1b, c). Canonical SREBP2 targets (regulating primarily cholesterol synthesis and uptake) were less affected by DGAT1/2 inhibition (Extended Data Fig. 1d). In adipocytes and U2OS cells, blocking TG synthesis can lead to activation of the unfolded protein response (UPR) or the ISR^8,36^. However, in our RNA-seq experiments, we did not detect robust changes in targets of the ISR or UPR upon DGAT inhibition, either under basal conditions or with oleate addition (Fig. 1h and Extended Data Fig. 1e).

### Impairing TG synthesis induces PPARα-mediated up-regulation of genes to increase fatty acid oxidation

To further test whether DGAT inhibition activates PPAR transcription factors in cells, we independently validated changes of well-known PPARα target genes by quantitative reverse transcription-PCR. Consistent with the RNA-seq analysis, these experiments showed two- to threefold up-regulation of *PDK4*, *CYP1A1*, and *CPT1A*, particularly when both DGATs were inhibited (Fig. 2a).

**Figure 2.**
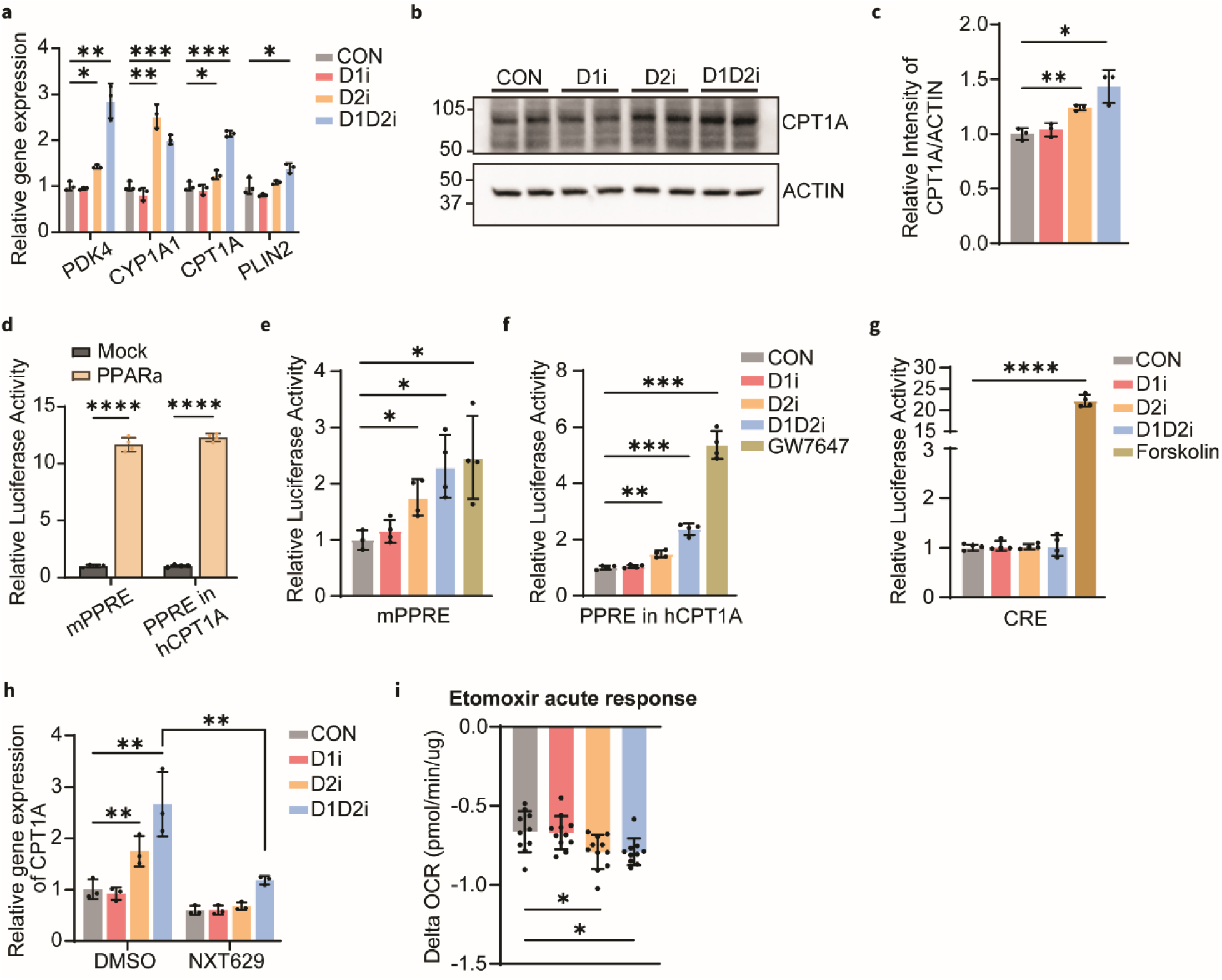
Inhibiting TG synthesis induces PPARα-mediated up-regulation of genes to increase fatty acid oxidation. **(a)** Bar graph showing relative mRNA expression of major upregulated PPARα target genes. Data are presented as mean ± SD (n = 3 biological replicates). ∗p < 0.05; ∗∗p < 0.01; ∗∗∗p < 0.001. **(b-c)** Western blot analysis (**b**) and quantification (**c**) showing increased CPT1A protein levels after DGAT2 inhibition and combined DGAT1 and DGAT2 inhibition. Data are presented as mean ± SD in bar graph (n = 3 for each treatment). ∗p < 0.05; ∗∗p < 0.01. **(d)** Dual-luciferase reporter assay analysis showing increased transcriptional activity of the mouse PPRE (mPPRE) and the human CPT1A PPRE upon human PPARα overexpression. Mean ± SD, n = 4 for each treatment, ∗∗∗∗p < 0.0001. **(e-f)** Bar graphs showing relative luciferase activity of the mouse PPRE (mPPRE, **e**) and the human CPT1A PPRE (PPRE in hCPT1A, **f**) in Huh-7 cells treated with DMSO (CON), DGAT1 inhibitor (D1i), DGAT2 inhibitor (D2i), or combined DGAT1 and DGAT2 inhibition (D1D2i) for 16 h. Huh-7 cells treated with GW7647, a PPARα agonist, were used as a positive control. Mean ± SD, n = 4. ∗p < 0.05; ∗∗p < 0.01; ∗∗∗p < 0.001. **(g)** Relative luciferase analysis of cAMP response element (CRE)-driven transcriptional activity after DGAT inhibitor treatment. Forskolin treatment served as a positive control. Mean ± SD, n = 4, ∗∗∗∗p < 0.0001. **(h)** Inhibition of PPARα prevents DGAT inhibition–induced CPT1A gene expression. Bar graph showing relative CPT1A mRNA expression after DGAT inhibition in the presence or absence of the PPARα antagonist NXT629. Error bars represent as mean ± SD of three biological replicates. Statistical significances were evaluated by two-way ANOVA, followed by Tukey-Kramer test. **p < 0.01. **(i)** Delta oxygen consumption rate (OCR) of Huh-7 cells treated with or without DGAT inhibition in response to etomoxir treatment. Values shown are the delta OCR after etomoxir injection. Mean ± SD, n = 10–12, ∗p < 0.05.

PPAR transcription factors are members of a family of proteins with some overlapping and some distinct transcriptional targets. *CPT1A* is a canonical PPARα target gene that catalyzes the conversion of acyl-CoAs to acyl-carnitines, a rate-limiting step in fatty acid oxidation. In agreement with the transcriptional changes, CPT1A protein levels were increased after DGAT2 inhibition and further increased with combined DGAT1 and 2 inhibition (Fig. 2b, c).

To evaluate transcriptional activity of PPAR after DGAT inhibition, we sought to identify a robust transcriptional readout and selected the PPAR reporter containing three repeat DR1 sites ^37^ or the peroxisome proliferator-activated receptors response elements (PPRE) within the human *CPT1A* promoter driving luciferase expression as reporter assays. Both PPREs were robustly activated by human PPARα overexpression (Fig. 2d). In contrast, overexpression of other human PPAR transcription factor family members, such as PPARβ or PPARγ, did not activate the reporter containing PPRE in human *CPT1A* promoter in Huh-7 cells (Extended Data Fig. 2). Inhibiting either DGAT2 or both DGAT1 and 2 induced PPRE reporter activity (Fig. 2e, f). In contrast, DGAT inhibition did not alter the activity of a cAMP response element (CRE) reporter, suggesting that the effect was specific to the upregulation of PPARα-mediated transcriptional activity (Fig. 2g). In addition, inhibiting PPARα with the compound NXT629 reversed the increase of *CPT1A* gene expression with DGAT inhibition (Fig. 2h), further suggesting that upregulation of genes due to DGAT inhibition is mediated by PPARα.

PPARα activation in general, and CPT1A induction specifically, increases fatty acid oxidation in different cellular and physiological contexts^38,39^. We therefore tested the effect of DGAT inhibition on fatty acid oxidation using Seahorse metabolic flux analysis. Cells were first treated with DGAT inhibitors and subsequently with etomoxir, an inhibitor of CPT1A. Cells showed decreased oxygen consumption rates (OCRs) in response to etomoxir, and this response was exacerbated in cells exposed to DGAT2 and combined DGAT inhibition, indicating higher levels of fatty acid oxidation (Fig. 2i).

### Impairing TG synthesis suppresses SREBP1-mediated gene expression to decrease fatty acid synthesis and desaturation

The strongest transcriptional effects of DGAT inhibition were on SREBP1 target genes. SREBP-transcription factors are synthesized as membrane-embedded precursors in the ER. Upon activation, they translocate from the ER to the Golgi apparatus where sequential proteolytic cleavages liberate an N-terminal, soluble protein fragment that enters the nucleus and activates transcription of target genes^26,40,41^. DGAT2 or combined DGAT1 and 2 inhibition for 16 h (or 4 h; Extended Data Fig. 3a, b) reduced the protein abundance of nuclear SREBP1 (SREBP1-N) (Fig. 3a). We also observed decreased levels of the membrane precursor (SREBP1-P), an effect even more pronounced with combined DGAT1 and 2 inhibition for 16 h (Fig. 3b). DGAT2 or combined DGAT1 and 2 inhibition reduced the ratio of nuclear to precursor SREBP-1 by 50% or 70%, respectively (Fig. 3c).

**Figure 3.**
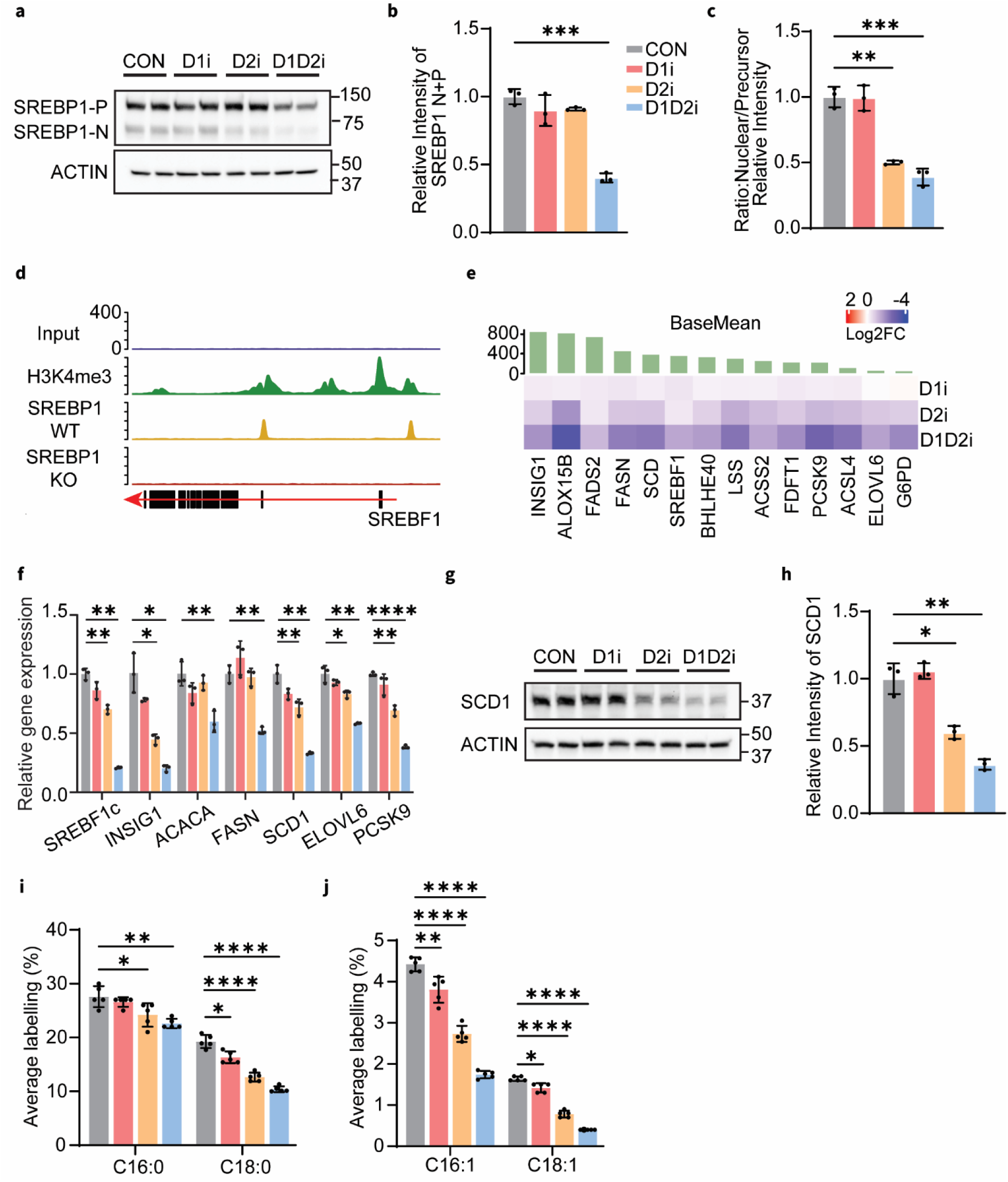
Inhibiting TG synthesis suppresses SREBP1-mediated expression of genes to decrease fatty acid synthesis. **(a)** Western blot of SREBP1 in whole-cell lysates from Huh-7 cells treated with DMSO (CON), DGAT1i inhibitor (D1i), DGAT2 inhibitor (D2i), or combined DGAT1 and DGAT2 inhibition (D1D2i) for 16 h. **(b-c)** Bar graphs showing quantification of relative total SREBP1 protein levels (SREBP1 N+P) (B) and the ratio of nuclear (N) to precursor (P) SREBP1 (C). Mean ± SD, n = 3, ∗∗p < 0.01; ∗∗∗p < 0.001. **(d)** Representative ChIP-seq tracks at the SREBF1 locus. Shown are input control, H3K4me3, and SREBP1 ChIP-seq signals in WT and SREBP1 KO Huh-7 cells. **(e)** Heatmap of SREBP1 ChIP-seq signals across target genes loci under DGAT inhibition. Shown are log2 fold-changes in SREBP1 binding peak intensity after DGAT inhibition, relative to control. The green bars represent the base mean of SREBP1 peak signal for each gene. **(f)** Relative gene expression of SREBP1 target genes performed by RT-qPCR. Huh-7 cells were treated with DGAT inhibitors for 16 h. **(g)** Western blot probing expression of SCD1 in whole-cell lysates of Huh-7 cells treated with DGAT inhibitors for 16 h. **(h)** Bar graphs showing quantification of relative SCD1 protein levels. Mean ± SD, n = 3, ∗p < 0.05; ∗∗p < 0.01. **(i-j)** *De novo* lipogenesis activity of saturated fatty acids (C16:0 and C18:0, **i**) and monounsaturated fatty acids (C16:0 and C18:0, **j**) in Huh-7 cells by DGAT inhibition. Cells were treated with DGAT inhibitors for 16 h, followed by incubation in deuterium-labeled medium for 6 h. Mean ± SD, n = 5, ∗p < 0.05; ∗∗p < 0.01; ∗∗∗∗p < 0.0001. Statistical significance was evaluated by unpaired two-tailed *t*-test.

We also examined the effects of DGAT2 or combined DGAT1 and 2 inhibition on SREBP2 activation. DGAT inhibition decreased SREBP2-N levels at 4 h, but these effects were not apparent at 16 h of treatment (Extended Data Fig. 3c-e). The few genes encoding enzymes within the cholesterol biosynthetic pathway that we found altered by DGAT2 or combined DGAT1 and 2 inhibition respond not only to SREBP2 but also SREBP1a^42,43^.

To directly investigate SREBP1 transcriptional activity towards its targets, we performed ChIP-seq of SREBP1 in Huh-7 cells treated with DGAT inhibitors for 16 h. As a specificity control for these experiments, we generated SREBP1 knockout (KO) cells and validated this cell line by western blot and SREBP1 gene expression (Extended Data Fig. 4a, b). ChIP-seq analyses of DNA bound by SREBP1-N showed specific enrichment of sequences from the *SREBF1* and *SCD1* promoters in WT, but not in SREBP1 KO cells (Fig. 3d and extended Data Fig.4c). DGAT2 or combined DGAT1 and 2 inhibition markedly decreased SREBP1 binding activity on target promoters, such as *SREBF1, INSIG1, ALOX15B, FASN* and *SCD1* (Fig. 3e).

The decreased binding of active, processed SREBP1-N was reflected in lower levels of mRNA produced from these genes. Combined DGAT1 and 2 inhibition repressed expression of many canonical SREBP1 target genes (e.g., *SREBF1, INSIG1, ACACA, FASN, SCD1, ELOVL6* or *PCSK9*), whereas DGAT2 inhibition alone had less dramatic and more selective effects (Fig. 3f). To test whether the effects of DGAT inhibition may be due to off-target effects of the inhibitors used, we analyzed mRNA levels after depleting either enzyme from cells. siRNA-mediated DGAT1 and 2 depletion resulted in a substantial and specific reduction in either DGAT enzyme (Extended Data Fig. 4d), as well as decreased levels of mature SREBP1-N (Extended Data Fig. 4e, f), and mRNA levels of its target genes (e.g., *SREBF1*, *FASN*, *SCD1* or *PCSK9*; Extended Data Fig.4g, h).

These changes also translated to altered protein levels in cells after DGAT inhibition. For instance, DGAT2 or combined DGAT1 and 2 inhibition resulted in decreased levels of the Δ9-desaturase SCD1, a key fatty acid desaturase and well-established SREBP1 target (Fig. 3g, h). Similarly, other protein targets, such as INSIG1, were reduced after DGAT2 or combined DGAT1 and 2 inhibition (Extended Data Fig. 4i). Since many of the genes regulated by SREBP1 function in *de novo* lipogenesis, we hypothesized that DGAT inhibition decreases the activity of this pathway. We performed metabolic tracing experiments with deuterated water (D₂O) as a stable isotope marker to measure fatty acid synthesis in Huh-7 cells treated with DGAT inhibitors for 16 h. Both DGAT2 and combined DGAT1 and 2 inhibition markedly decreased the percentage of ^2^D incorporation into palmitate and stearate, as well as into accompanying monounsaturated fatty acids (16:1 and 18:1), indicating reduced *de novo* lipogenesis (Fig. 3i, j).

### Impairing TG synthesis independently impairs SREBP1 precursor synthesis and proteolytic processing

To test whether the changes in abundance of SREBP1 precursor and nuclear forms are independent or due to feed-forward regulation of its own gene promoter, we sought to experimentally separate these effects of DGAT inhibition. To determine if changes of SREBP precursors are due to transcriptional control, we expressed SREBP1 from a heterologous thymidine kinase promoter in SREBP1 KO cells and analyzed the effects of DGAT inhibition. Without its endogenous promoter, DGAT2 or combined DGAT1 and 2 inhibition did not decrease SREBP1 precursor levels, but still decreased levels of cleaved SREBP1-N (Fig. 4a-c), revealing distinct effects of DGAT inhibition on SREBP1 mRNA expression and precursor protein processing.

**Figure 4.**
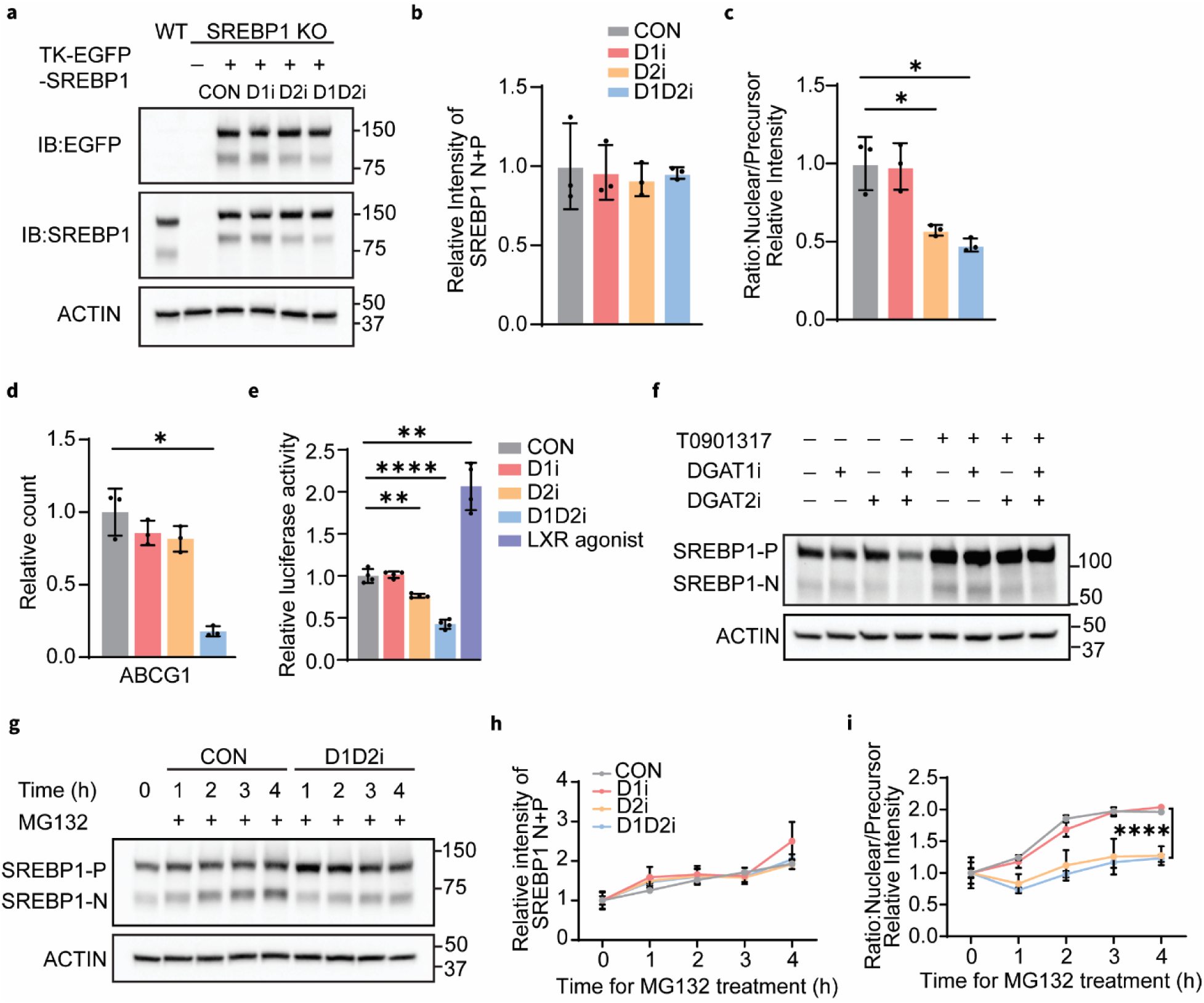
Inhibiting TG synthesis independently impairs SREBP1 precursor synthesis and proteolytic processing. **(a)** Decreased nuclear SREBP1 levels are independent of transcriptional regulation. SREBP1 KO Huh-7 cells transiently expressing TK-EGFP–SREBP1 were treated with DGAT inhibitors for 16 h. Western blot analysis of whole-cell lysates from WT Huh-7 cells and treated SREBP1 KO cells was performed using anti-EGFP and anti-SREBP1 antibodies. **(b-c)** Bar graphs showing quantification of relative total SREBP1 protein levels (SREBP1 N+P; **b**) and the ratio of nuclear (N) to precursor (P) SREBP1 (**c**). Mean ± SD, n = 3, ∗p < 0.05. Statistical significance was evaluated by unpaired two-tailed *t*-test. **(d)** Relative counts of *ABCG1* in Huh-7 cells derived from RNA-seq analysis. *p < 0.05. **(e)** Relative luciferase analysis of LXR response element (LXRE)–driven transcriptional activity after DGAT inhibitor treatments. T0901317 (LXR agonist) treatment served as a positive control. Mean ± SD, n = 4, ∗∗p < 0.01; ∗∗∗∗p < 0.0001. **(f)** Western blot analysis of SREBP1 in whole-cell lysates of Huh-7 cells treated with DGAT inhibitors in the presence or absence of T0901317. **(g)** Western blot analysis of SREBP1 in whole-cell lysates of Huh-7 cells treated with DMSO (CON) or combined DGAT1 and DGAT2 inhibitors (D1D2i) in the presence of MG132 at the indicated time points. **(h-i)** Quantification of relative total SREBP1 protein levels (SREBP1 N+P; **h**) and the ratio of nuclear (N) to precursor (P) SREBP1 (**i**) at the indicated time points. Mean ± SD, n = 3, ∗∗∗∗p < 0.0001. Statistical significance was calculated by two-way ANOVA with repeated measurements.

We hypothesized that the transcriptional effects of DGAT inhibition on SREBP1 were mediated by LXR, a transcription factor that regulates SREBP1^44^. We therefore tested the effects of DGAT inhibition on LXR activity by examining levels of LXR target gene transcripts in our RNA-seq data. The canonical LXR target gene *ABCG1* was significantly decreased after combined DGAT1 and 2 inhibition (Fig. 4d). Consistent with these findings, the activity of a LXRE luciferase reporter was diminished with DGAT2 or combined DGAT1 and 2 inhibition (Fig. 4e). Conversely, activation of LXR by agonists abolished the decrease in SREBP1 precursor levels in DGAT inhibition but did not block effects on SREBP1 mature levels (Fig. 4f). Thus, one effect of DGAT inhibition appears to be the downregulation of LXR-mediated SREBP1 expression.

We next investigated how the processed form of SREBP1 is decreased. First, we distinguished between decreased cleavage of the precursor or accelerated degradation of the processed SREBP1. SREBP1-N is degraded by the proteasome^45^. We measured SREBP1 levels during DGAT inhibition after proteasome inhibition by MG132. Under these conditions, MG132 blocked protein degradation because total levels of SREBP1 precursor and the processed form increased, but DGAT2 or combined DGAT1 and 2 inhibition still reduced SREBP1-N levels (Fig. 4g-i and Extended Data Fig. 4j, k).

Together, these data suggest two independent effects mediate the decreased SREBP response during DGAT inhibition: an LXR-mediated downregulation of *SREBP1* transcription and a decrease in SREBP1-processing to the mature, transcriptionally active form.

### Impairing TG synthesis results in many changes of membrane lipid composition

How SREBP1 is regulated has been a long-standing mystery, and we reasoned DGAT inhibition may provide an ideal setting to answer this question. We therefore sought to determine how DGAT inhibition regulates SREBP1 cleavage, the key step in its regulation. Since DGAT enzymes catalyze reactions of lipid metabolism, we hypothesized that changes in cellular lipids would mediate the effects on SREBP1. Consistent with this idea, a previous report concluded that changes in ER levels of PE mediate changes in SREBP1 activity^25^.

To test this and other possible lipid mediators of the response in an unbiased manner, we performed comprehensive and quantitative lipidome analyses of cells with DGAT inhibition under various conditions. These included different time points (4 and 16 h), as well as analysis of ER microsomes.

As expected, combined DGAT1 and 2 inhibition in Huh-7 cells, and to a lesser extent DGAT2 inhibition, reduced the abundance of TG, the product of their reactions, after 16 h of treatment (Fig. 5a). Similarly, for both DGAT2 and combined DGAT1 and 2 inhibition, we detected accumulation of diacylglycerol (DG) with increases in several free fatty acids, such as 20:4 and 22:6, whereas levels of free 16:0, 18:1, and 18:2 were unchanged (Fig. 5b, c and Extended Data Fig. 5a). Since cholesterol mediates SREBP regulation and because its level is not always accurately measured in mass spectrometry, we quantified cholesterol levels in whole cell and microsome fractions using a fluorometric assay and found no change in cholesterol levels after 16 h of DGAT inhibitor treatment in Huh-7 cells (Fig. 5d).

**Figure 5.**
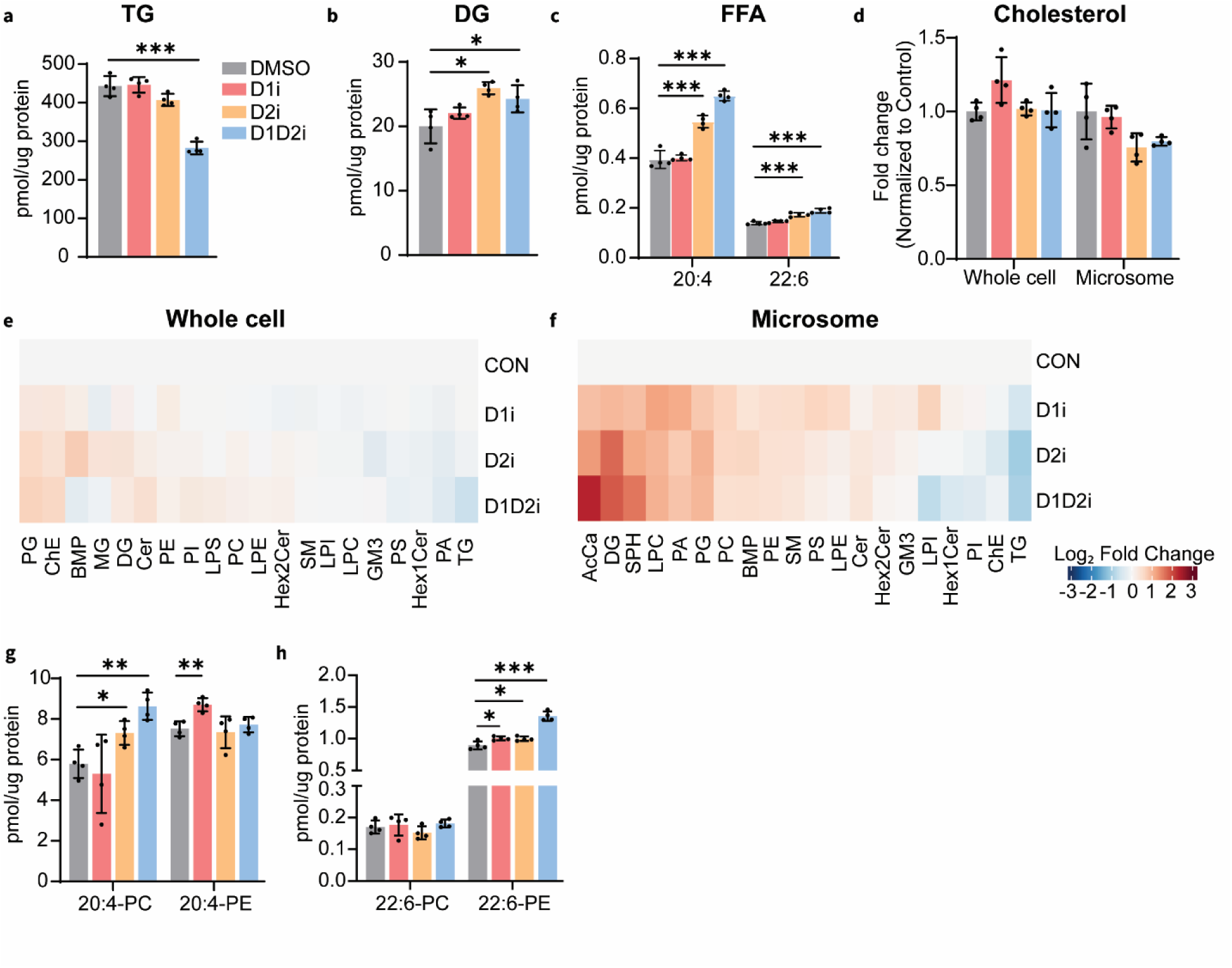
Inhibiting TG synthesis changes membrane lipid composition. **(a)** TG levels in Huh-7 cells treated with DMSO (CON), DGAT1i inhibitor (D1i), DGAT2 inhibitor (D2i), or combined DGAT1 and DGAT2 inhibition (D1D2i) for 16 h. ∗∗∗p < 0.001. **(b)** Diacylglycerol (DG) levels in Huh-7 cells treated with DGAT inhibitors for 16 h. ∗p < 0.05. **(c)** Free fatty acid levels in Huh-7 cells treated with DGAT inhibitors for 16 h. ∗∗∗p < 0.001. **(d)** Cholesterol levels in whole-cell lysis and microsome of Huh-7 cells treated with DGAT inhibitors for 16 h. **(e)** Heatmap analysis of fold-change of major lipid class abundances in whole-cell lysates of Huh-7 cells treated with DGAT inhibitors for 16 h, normalized to control. Data plotted as mean of the log2 (fold-change). **(f)** Heatmap analysis of fold-change of major lipid class abundances in microsomes of Huh-7 cells treated with DGAT inhibitors for 16 h, normalized to control. **(g-h)** Bar graphs showing the abundances of phospholipids containing arachidonic acid (20:4; **g**) or docosahexaenoic acid (22:6; **h**) in whole-cell lysates of Huh-7 cells treated with DGAT inhibitors for 16 h. Statistical significance was evaluated by unpaired two-tailed *t*-test. Mean ± SD, n = 4, ∗p < 0.05; ∗∗p < 0.01; ∗∗∗p < 0.001.

Regulation of SREBP processing presumably occurs at ER or Golgi membranes. Although these organelles represent the major pool of phospholipids in cells, regulatory changes due to DGAT inhibition may be masked by lipids in other organelles^25^. To test this possibility, we analyzed lipid changes in ER and Golgi microsomes. Similarly to whole cell lipids, we observed an increase in DGAT substrates, such as DGs, and a decrease in products, such as TGs (Fig. 5e, f). However, we observed no significant changes in levels of PE or phosphatidylcholine (Extended Data Fig. 5b, c).

DG is an important second messenger that activates protein kinases and regulates numerous cellular responses. DG levels increased in multiple tested conditions, such as 4 h inhibitor treatment or OA supplement (Extended Data Fig. 5d). However, analysis of lipid changes in cells with siRNA-mediated DGAT2 depletion led to suppression of SREBP1 activity in the absence of DG changes (Extended Data Fig. 5e).

Fatty acids, and specifically PUFAs, are sufficient to suppress SREBP1 activity in HEK293 cells^22^. In some experiments, we detected increased levels of unesterified 20:4 and 22:6 fatty acids. Yet, despite this increase of PUFAs and PUFA-containing phospholipids in some conditions (e.g., in DGAT inhibition for 16 h; Fig. 5g, h), similar changes were not found in other conditions where SRBEP1 suppression is observed (e.g., DGAT inhibition under OA supplement; Extended Data Fig. 5f).

To test which of the lipid changes corresponds to changes observed with genetic ablation of DGAT2 in murine hepatocytes that leads to SREBP1 suppression, we also determined lipid changes for mouse liver. Consistent with the changes in cells, loss of DGAT2 in liver resulted in reductions in hepatic TG levels (Extended Data Fig. 5g). Unlike the increased DG and PUFAs levels in cells, free 20:4 and 20:6 levels were unchanged, but free 18:1 fatty acid and DG levels were decreased in liver (Extended Data Fig. 5g, h).

Thus, by examining multiple conditions in cells or liver where SREBP1 activation was suppressed, we found no clear lipid change that consistently correlated with this response.

### Alterations in ER membrane fluidity regulate SREBP1 processing

Failure to detect a single lipid species or class with levels correlating to changes in SREBP1 activity led us to consider the alternative hypothesis that changes in the collective properties of the ER membrane mediate the response to DGAT inhibition. Specifically, we hypothesized that DGAT inhibition results in a variety of lipid changes (e.g., DG, PUFA, PE) that increase the fluidity of the ER and affect SREBP1 processing.

To directly test this model, we measured changes in ER membrane fluidity across different conditions. For this, we stained cells with ER-laurdan, a dye that accumulates in the ER and that changes its fluorescence behavior depending on water accessibility to the fluorophore^46^. This is a proxy readout for alterations in membrane fluidity and can be quantitated as a generalized polarization (GP) index for individual cells. Treatment with DGAT2 or combined DGAT1 and 2 inhibitors for 4 h decreased the GP index of ER membranes, indicating increased membrane fluidity (Fig. 6a, b). This reduction in GP index was also evident after 16 h of DGAT2 or combined DGAT1 and 2 inhibition (Extended Data Fig. 6a). To test whether the increase in membrane fluidity was reflected in a change in diffusion rate of an ER-localized protein, we expressed the transmembrane segment of cytochrome b5 (CYTB5) fused to a Halo tag in Huh-7 cells and tracked single molecules using minimal fluorescence photon fluxes (MINFLUX) nanoscopy (Fig. 6c). Combined DGAT1 and 2 inhibition increased the apparent diffusion of ER-localized CYTB5 by ∼20% (Fig. 6d), in agreement with an increase in membrane fluidity.

**Figure 6.**
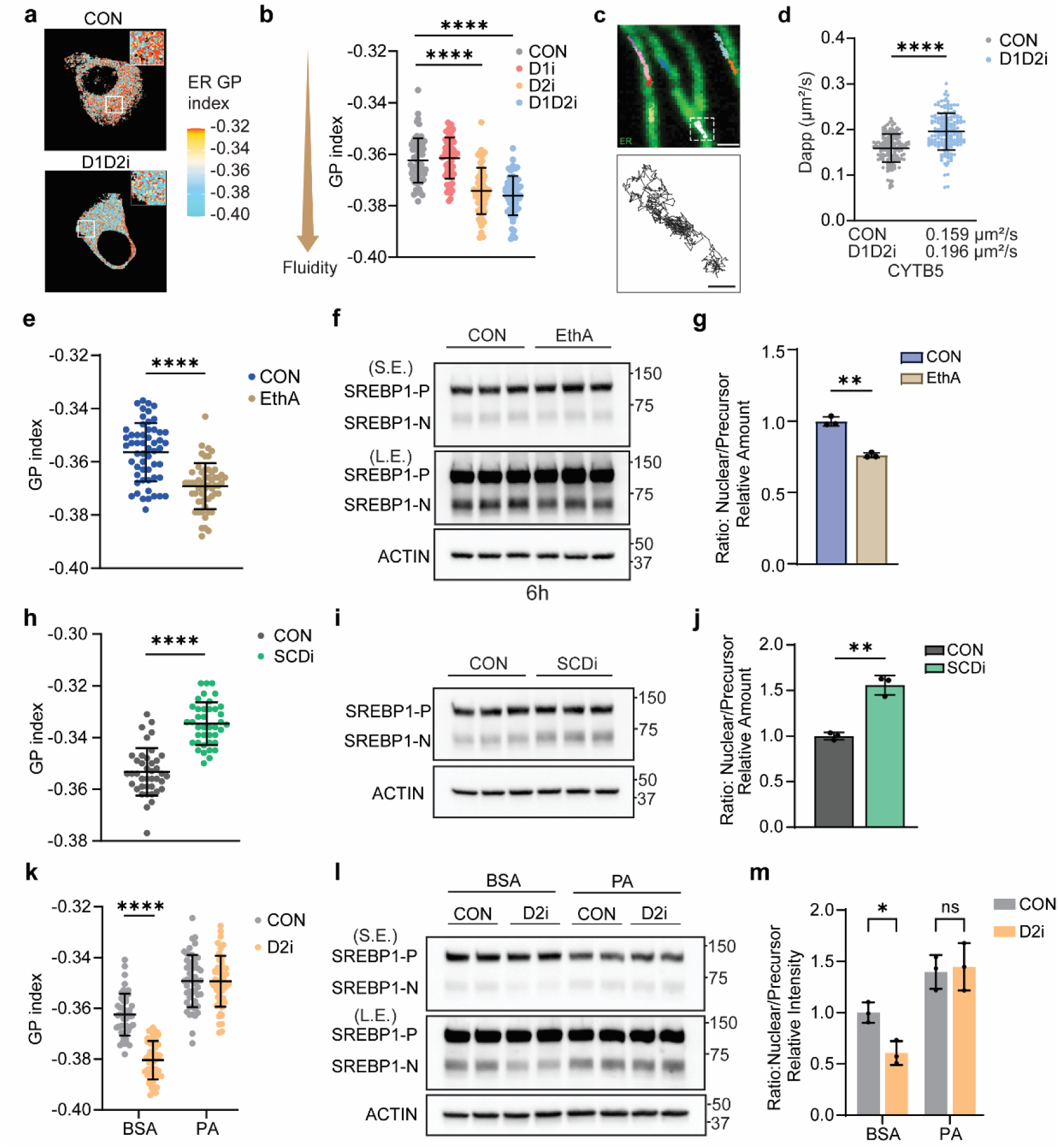
Changes in ER membrane fluidity correlate with altered SREBP1 processing. **(a)** ER-Laurdan generalized polarization (GP) maps of Huh-7 cells under control (CON) and combined DGAT1 and DGAT2 inhibition (D1D2i). Huh-7 cells treated with DGAT inhibitors for 4 h. **(b)** Quantification of GP index of Huh-7 treated with DMSO or DGAT inhibitors for 4 h. Mean ± SD, n = 60 cells, ∗∗∗∗p < 0.0001. Statistical significance was evaluated by unpaired two-tailed t-test. **(c)** MINFLUX single-molecule trajectories of truncated-CYTB5. Scale bars, 500 nm (up) and 100 nm (down). **(d)** Diffusion coefficients analysis for truncated-CYTB5 in Huh-7 cells treated with DMSO(CON) or both DGAT inhibitors(D1D2i) for 16 h, calculated using a 5-ms rolling-window mean squared displacement (MSD) analysis. Each dot represents one track, N(Control)=157, N(D1D2i) =162. **(e)** Quantification of ER GP index in Huh-7 cells treated with ethanolamine (EthA, 100 μM) for 6 h. Data are presented as mean ± SD (n = 55 cells). ∗∗∗∗p < 0.0001, determined by unpaired two-tailed *t*-test. **(f-g)** Western blot analysis and quantification of SREBP1 in whole-cell lysates. Huh-7 cells were treated with ethanolamine for 6 h. Mean ± SD, n = 3, ∗∗p < 0.01, unpaired two-tailed *t*-test. **(h)** Quantification of ER GP index in Huh-7 cells treated with SCD1 inhibitor (MK-8245, SCDi) for 16 h. Data are presented as mean ± SD (n = 55 cells). ∗∗∗∗p < 0.0001, determined by unpaired two-tailed *t*-test. **(i-j)** Western blot analysis and quantification of SREBP1 in whole-cell lysates. Huh-7 cells were treated with SCD1 inhibitor for 16 h. Mean ± SD, n = 3, ∗∗p < 0.01, unpaired two-tailed *t*-test. **(k)** Quantification of ER GP index in Huh-7 cells pretreated with palmitate (PA, 200 µM) for 4 h and subsequently treated with DMSO or DGAT2 inhibitor in the presence of BSA or PA for 4 h. Data are presented as mean ± SD (n = 60 cells). ∗∗∗∗p < 0.0001, determined by unpaired two-tailed *t*-test. **(l-m)** Western blot analysis and quantification of SREBP1 in whole-cell lysates. Huh-7 cells were pretreated with BSA or 200 μM PA for 4 h and then treated with DMSO or DGAT2 inhibitor in the presence BSA or PA. Mean ± SD, n = 3, ∗p < 0.05, Statistical significance was determined by two-way ANOVA, followed by Šídák’s multiple comparisons test for selected pairwise comparisons.

We next asked whether regulation of SREBP1 by altered ER membrane properties represents a more general mechanism across conditions that modulate SREBP1 activity. Elevated PE levels are associated with reduced SREBP1 cleavage^25^. We therefore tested whether ethanolamine supplementation to increase PE levels in cell membranes is sufficient to increase membrane fluidity and decrease SREBP1 processing. As expected, ethanolamine supplementation increased PE levels and decreased levels of DG, a PE-synthesis substrate (Extended Data Fig. 6b). Ethanolamine supplementation of cell growth media also decreased the ER-laurdan GP index, indicating increased ER membrane fluidity (Fig. 6e). In agreement with our model, ethanolamine supplementation also decreased SREBP1 cleavage (Fig. 6f, g).

We next asked whether the SREBP system responds to decreased ER membrane fluidity by enhancing SREBP1 processing. To test this hypothesis, we inhibited SCD1, which decreased the levels of mono-unsaturated fatty acids in glycerolipids (Extended Data Fig. 6c). Consequently, this reduced ER membrane fluidity, as evident from an increased GP index of ER-laurdan (Fig. 6h), and this correlated with increased SREBP1 processing (Fig. 6i, j).

Finally, to determine if increased membrane fluidity is required for inhibiting SREBP1 processing in the setting of DGAT inhibition, we used palmitate (C16:0) to stiffen the ER membrane and test if this could reverse the effects of DGAT inhibition on SREBP1 activation. With palmitate supplementation to the medium, the GP index of Huh-7 cells increased, indicating that palmitate addition was sufficient to decrease ER membrane fluidity (Extended Data Fig. 6d). Palmitate supplementation also reversed the DGAT2 inhibition–induced reduction in GP index, indicating reversal of membrane fluidity changes (Fig. 6k). Consistently, palmitate supplementation of cell growth medium restored the reduced levels of nuclear SREBP1 caused by DGAT2 inhibition (Fig. 6l, m). Stearic acid (SA, C18:0) supplementation produced a similar restorative effect (Extended Data Fig. 6e, f). Together, these perturbations establish a correlation between ER membrane fluidity and SREBP1 activation.

## Discussion

Here we identified a homeostatic response that is engaged when fatty acid flux exceeds the capacity for TG synthesis and storage in hepatocytes. Under these conditions, cells coordinately activate fatty acid oxidation and suppress SREBP1-mediated lipogenesis, thereby reducing lipotoxic stress. These findings uncover a lipid-homeostatic system that couples TG synthesis capacity to the regulation of fatty acid metabolism. Mechanistically, SREBP1 cleavage-activation responds when ER membrane fluidity changes, suggesting that SREBP1 functions as part of a membrane-homeostatic circuit that adjusts lipid synthesis and desaturation to maintain membrane properties.

PPARα activation with DGAT inhibition likely reflects transient accumulation of PUFAs, particularly arachidonic acid (20:4) and docosahexaenoic acid (22:6), reported activators of PPARs^47,48^. In Huh-7 cells, this response was most robust with DGAT2 or combined DGAT1/2 inhibition, indicating a distinct function of DGAT2 in these cells. However, the relative contributions of DGAT1 and DGAT2 to this response may differ, consistent with the tissue-specific functions of these enzymes^49,50^. Inasmuch as no PPARα response was observed when DGAT2 was deleted in murine liver, fatty acids may be metabolized on longer timescales, thus blunting this acute response to DGAT inhibition. In agreement with this notion, short-term knockdown of DGATs or injection of DGAT inhibitors in mice increased *CPT1A* gene expression in liver^19,51^. Moreover, DGAT1 and 2 deletion in white or brown adipocytes upregulates fatty acid oxidation^52–54^. Together, these findings support a model in which unbuffered fatty acids are acutely redirected toward oxidation when TG-storage capacity is exceeded.

The strongest transcriptional response to DGAT inhibition was suppression of SREBP1 target genes. DGAT2 inhibition or combined DGAT1/2 inhibition resulted in sustained downregulation of lipogenic gene expression. Specifically, expression of SREBP1 target genes involved in unsaturated fatty acid synthesis (e.g., *SCD1*, *FADS1* and *FADS2*) was consistently decreased in Huh-7 cells treated with DGAT inhibitors and in livers from Liv-DGAT2 KO mice. These findings are consistent with reports on the lipogenic gene expression consequences of DGAT2 depletion or knockdown in liver or white adipose tissue, both of which exhibit high levels of *de novo* lipogenesis ^19,21,50^. In addition, PNPLA3 expression was reduced in both mouse liver and human cells. Since PNPLA3 may preferentially hydrolyze PUFA-containing TGs^55^, its regulation by SREBP1 may further contribute to controlling membrane lipid composition and desaturation. DGAT inhibition reduced SREBP1 activation both by decreasing expression of the precursor protein and by suppressing cleavage activation, the key regulatory step controlling SREBP transcription factor activity^56,57^. Reduced precursor synthesis appears to be linked to diminished LXR activity^58^, as DGAT inhibition decreased LXR DNA-binding activity and corresponding expression of LXR target genes, such as *ABCG1*, whereas LXR agonism restored SREBP1 precursor levels.

How SREBP1 cleavage is regulated has been a long-standing mystery. Previous studies implicated changes in the levels of multiple lipid species, including PUFAs^22^, PE^59^, and PC^60,61^ in regulation of SREBP1 processing. Specifically, for DGAT2 inhibition, increased PE levels have been reported to suppress SREBP1 processing in mammalian cells^25^. However, while some of these lipid changes may be sufficient to alter SREBP1 processing, across multiple experimental systems, we did not identify any lipid species that consistently correlated with suppression of SREBP1 activation in response to DGAT inhibition. For instance, we did not observe consistent alterations in PE levels in either whole-cell lysates or microsomal fractions under conditions in which we observed SREBP1 suppression. Similarly, DGAT2 inhibition increased levels of PUFAs and DG in Huh-7 cells, but murine liver lacking DGAT2 showed SREBP1 suppression without changes in these lipids.

Instead, SREBP1 cleavage and activation occurred with changes in ER membrane fluidity. DGAT inhibition increased ER membrane fluidity and suppressed SREBP1 activation, whereas saturated fatty acids exerted opposite effects on membrane properties and SREBP1 processing. This effect was also found when we perturbed ER membrane fluidity by challenging cells with incubation of other lipids or precursors. Control of SREBP1 by ER membrane properties is consistent with the suppression of SREBP1 in livers of PUFA-rich diet–fed mice, where increased docosahexaenoic acid–containing phospholipids are associated with elevated membrane fluidity^62^. Together, a model emerges in which SREBP1 responds to ER membrane properties to maintain lipid homeostasis. In this model, reduced membrane fluidity promotes SREBP1 activation, increasing fatty acid synthesis and unsaturation to restore fluidity, whereas excess fatty acid accumulation increases membrane fluidity and suppresses SREBP1 activation to reduce fatty acid synthesis. This model resembles membrane homeostatic systems in other organisms, such as yeast, where Mga2 senses membrane changes to regulate expression of the fatty acid Δ-9 desaturase Ole1^63,64^.

In mammal cells, homeostatic control of ER membrane properties is essential for supporting efficient cargo trafficking through the secretory pathway^65,66^. How the SREBP1 system senses changes in membrane properties remains unclear molecularly. For SREBP2, its response to cholesterol has been linked to conformational changes of SCAP, a chaperone for SREBP trafficking out of the ER, essential for cleavage by proteases of the Golgi apparatus^56,67^. Intriguingly, changes in SREBP2 activation in response to changes of cholesterol occur with high cooperativity, most consistent with a change in membrane properties at a threshold, analogous to a membrane phase transition^68–70^. Some lipid changes observed with DGAT inhibition, such as increased microsomal DG, may contribute to SREBP1 regulation through additional mechanisms, including protein phosphorylation^24^.

Our findings may have relevance for understanding the detrimental effects of diets rich in saturated fatty acids^71,72^, which appear to also induce the membrane homeostatic response activating the SREBP1 pathway reported here. They may also have direct relevance for therapies targeting DGAT2 in metabolic dysfunction-associated steatotic liver disease/metabolic dysfunction-associated steatohepatitis, where *de novo* lipogenesis is a contributor^73,74^. DGAT2 inhibition reduces hepatic lipid accumulation and improves liver steatosis in animal models and clinical studies^75–78^, and our findings suggest these effects are achieved not only through reduced TG synthesis but also broader adaptive responses affecting fatty acid oxidation and lipogenesis. Our studies now shed light on how these effects are achieved and suggest that exposure to dietary fatty acids, especially saturated ones, may be variables to consider in future analyses.

## Acknowledgments

We thank Drs. Morris Birnbaum, Robert Dullea and Kendra Bence, as well as members of the Farese & Walther laboratory for helpful discussions. Dr. Niklas Mejhert and Aditi Jatkar for initial discussions and experiments. Dr. Itay Budin for ER-laurdan reagent, Dr. Christina Pyrgaki (MSKCC advanced light microscopy innovation laboratory) for assistance with microscopy, Dr. Ritchie Ly (MSKCC lipidomics innovation laboratory) for lipidomics assistance, and Gary Howard for editorial assistance. This work was supported by NIH R01 DK101579 (T.C.W and R.V.F) and R35GM158422 (T.C.W). We acknowledge NIH/NCI Cancer Center Support Grant (P30 CA008748) at MSKCC and the Center for Causes and Prevention of Cardiovascular Disease (CAP-CVD) at the Harvard Chan School of Public Health (S.H.). T.C.W. is a Howard Hughes Medical Institute investigator.

## Author contributions

X.X., R.V.F., and T.C.W. conceived the project, and R.V.F. and T.C.W. acquired project funding. X.X. performed most of the experiments. Y.A.A. helped to perform and analyze general lipidomic and free fatty acid profiles. S.H. and O.T. performed *de novo* lipogenesis assays. W.T. helped to perform and analyze the ER-laurdan assay. A.M. helped with analysis of MINFLUX data. Q.Z. helped with the reporter assay. L.C.A. helped with MINFLUX experiments. X.X., R.V.F., and T.C.W. wrote and all authors edited the manuscript.

## Declaration of interests

The authors have declared that no conflict of interest exists.

## Methods

### Animal study

Animal procedures were approved by the Institutional Animal Care and Use Committee of the Memorial Sloan Kettering Cancer Centre on animals, following NIH guidelines. Liver-specific *Dgat2* knockout mice were generated by crossing *Dgat2^flox/flox^*mice of C57BL/6J background with mice expressing Cre recombinase under control of the albumin promotor (B6.Cg-Tg(Alb-Cre)21MGN/J; The Jackson Laboratory, Bar Harbor, ME), as described^21^. This caused deletion of exons 3–4 in *Dgat2* and more than 80% reduction in hepatic *Dgat2* transcript levels. All mice were maintained in a pathogen-free barrier facility with a 12 h light/12 h dark cycle and *ad libitum* access to water and food. Equal number of male mice (8–10 weeks old) were used for experiments.

### Cell culture

The human hepatoma cell line Huh-7 (JCRB, JCRB0403) was obtained from JCRB cell bank and cultured in Roswell Park Memorial Institute (RPMI) 1640 medium (Thermo Fisher Scientific, 61870127) supplemented with 10% heat-inactivated FBS and penicillin/streptomycin at 37°C in 5% CO2.

### Preparation of albumin-bound fatty acids stock

A 10 mM oleic acid (OA) (Millipore-Sigma, O1383) stock solution was prepared by slowly adding OA to essentially fatty acid free–bovine serum albumin (BSA) (Millipore-Sigma, A6003), followed by incubation at 37°C with continuous shaking for at least 1 h to facilitate complete solubilization prior to cell treatment.

Palmitate (Millipore-Sigma, P9767) was prepared by dissolving sodium palmitate in water and melting the solution in a boiled water bath for a few minutes. A 10 mM palmitate-BSA conjugate was generated by adding the warm palmitate solution to pre-warmed BSA and incubating the mixture at 42°C for 30 min, filtered through a 0.22-μm filter, aliquoted and stored in −20°C.

### Lipid extraction and thin-layer chromatography

Effects of DGAT inhibition on TG synthesis in Huh-7 cells was determined by measuring the incorporation of [^14^C]-OA (50 μCi/μmol) (American Radiolabeled Chemicals, Inc, ARC 0297) into TG. Cells were seeded into six-well plates and cultured overnight. Cells were labeled with [^14^C]-OA in presence of DGAT inhibitors (DGAT1i, PF-04620110, Millipore-Sigma, PZ0207; DGAT2i, PF-06424439, Millipore-Sigma, PZ0233) for 16 h. After labeling, cells were washed three times with ice-cold PBS. Lipids were extracted directly from the wells using a hexane:isopropanol (3:2, v:v) mixture with gentle shaking for 10 min at room temperature. The extraction was repeated twice to ensure complete lipid recovery. The combined extracts were dried under nitrogen stream and separated by thin-layer chromatography (TLC) using a solvent system of hexane:diethyl ether:acetic acid (80:20:1, v:v:v). TLC plates were exposed to phosphoimaging cassettes overnight and visualized using an Amersham Typhoon biomolecular imager. Lipid standards on the same TLC plates were subsequently stained with iodine vapor for band identification.

### Plasmid construction

The following plasmids were kind gifts: LXRE_Luc from Thomas Burris (Addgene, 177622) and PPRE X3-TK-luc from Bruce Spiegelman (Addgene, 1015). The pcDNA3.1-PPAR isoforms plasmids were purchased from GenScript (pcDNA3.1-C-human PPARA, OHu20099D; pcDNA3.1-C-human PPARD, OHu20193D; pcDNA3.1-C-human PPARG, OHu24140D). pGL4.29[luc2P/CRE/Hygro] and pRL-TK vectors were purchased from Promega E8471 and E2241).

The luciferase reporter containing the PPRE from the human *CPT1A* promoter (PPRE in hCPT1A) was generated by inserting annealed oligonucleotides into a restriction enzyme–digested pGL4.29 vector. For expression of full length of SREBP1, the common coding region shared by SREBP1a and SREBP1c was amplified using Phusion High-Fidelity PCR Master Mix with HF Buffer (New England Biolabs, M0531S) and cloned into the pTK-EGFP vector. A truncated CYTB5 fragment was amplified from pLVX-EF1a-EGFP-CYTB5 (Addgene 134869) and cloned into the pTK-Halo vector using a HiFi DNA assembly kit (New England Biolabs). All restriction enzymes were from New England Biolabs.

### RNA sequencing and analysis

Huh-7 cells were seeded in six-well plates and cultured overnight. Adherent cells were washed once with pre-warmed PBS and then treated with RPMI containing lipid-depleted serum (Omega Scientific, FB-50) supplemented with DMSO, 10 µM DGAT1 inhibitor, 10 µM DGAT2 inhibitor, or both. After 16 h of treatment, cells were lysed, and total RNA was isolated using the RNeasy Kit (Qiagen, 74106).

Library construction and sequencing were performed by the MSKCC Integrated Genomics Operation Core. The RNA-seq pipeline was performed by the bioinformatics core at MSKCC. Briefly, after RiboGreen quantification and quality control by an Agilent BioAnalyzer, 500 ng of total RNA with RIN values of 10 underwent polyA selection and library preparation using the TruSeq Stranded mRNA LT Kit (Illumina). Samples were run on a NovaSeq 6000 in a PE100 run, using the NovaSeq 6000 S4 Reagent Kit (200 Cycles) (Illumina). An average of 20 million paired reads were generated per sample. Ribosomal reads represented 0.07–0.14% of the total reads generated and the percent of mRNA bases averaged 93%. FASTQ files were mapped to the target genome using the STAR aligner^79^, which performs genomic alignment and resolves reads spanning splice junctions. Resulting SAM files were processed with Picard tools to add read groups and convert to sorted BAM format. Gene-level read counts were generated using HTSeq, and raw count matrices were normalized and analyzed for differential expression using the DESeq2 package in R/Bioconductor.

### RNA extraction and Quantitative Real-Time PCR (qRT-PCR)

Total RNA was extracted from cells treated with DGAT inhibitors and liver tissues using the RNeasy Kit (Qiagen) and the manufacturer’s instructions. RNA quality and concentration were evaluated using a NanoDrop (Thermo Scientific), and equal amounts of RNA were used for downstream applications. Complementary DNA (cDNA) was synthesized using an iScript cDNA synthesis kit (BioRad, 1708891). qRT-PCR was performed with Power SYBR Green PCR Master Mix (Applied Biosystems™, 4367659) and amplification curves were recorded on a Bio-Rad CFX1000 Touch thermal cycler. Primer sequences for all target genes are listed in Table S1. Gene expression was normalized to the housekeeping gene cyclophilin and quantified using the delta-delta Ct method^80^.

### SDS-PAGE and western blot

Cells were seeded in 100-mm dishes and, after two washes, scraped into ice-cold PBS. Cells were pelleted at 700 × g for 5 min and resuspended in RIPA lysis buffer (Thermo Scientific, 89901) containing protease inhibitors for 30 min. Protein contents were measured with the BCA protein assay (Thermo Fisher Scientific, 23225). Normalized protein lysates were denatured in Laemmli sample buffer (Bio-Rad, 1610747) at 37°C for 30 min. Proteins were separated on 4–15% gradient SDS-PAGE gel and transferred to nitrocellulose membranes at 100 V for 1 h. Membranes were blocked in TBST supplemented with 5% Blotto, non-fat dry milk (Santa Cruz Biotechnology, sc-2325) at room temperature for 1 h and subsequently incubated in primary antibodies overnight at 4°C with gently shaking: anti-SREBP1 (1:1000, Millipore-Sigma, MABS1987), anti-SREBP2 (1:1000, Cell Signaling Technology, 25940), anti-ACTIN (1:1000, Santa Cruz Biotechnology, sc-47778), anti-CPT1A (1:1000, Cell Signaling Technology, 12252), anti-SCD1 (1:1000, Cell Signaling Technology, 2438), anti-CALNEXIN (1:1000, Santa Cruz Biotechnology, sc-46669), anti-PCNA (1:1000, Cell Signaling Technology, 13110), anti-GFP (1:1000, Abcam, ab290), anti-INSIG1 (1:1000, this paper). Membranes were washed three times with TBST for 5min and incubated with appropriate horseradish peroxidase–conjugated secondary antibodies (1:5000, Santa Cruz Biotechnology) at room temperature for 1 h. After another washing, membranes were developed using SuperSignal West Pico (Thermo Scientific, 34580) and imaged on a ChemiDoc XRS+ (BioRad).

### Transient transfection luciferase assay

Huh-7 cells were seeded in 24-well plates before transfection. Plasmid transfection was performed with FuGene HD transfection reagent (Promega, E2311), according to the manufacturer’s protocol. For each transfection reaction, the firefly luciferase reporter plasmid was co-transfected with the control plasmid pRL-TK encoding Renilla luciferase at 1:10 ratio. At 24 h after transfection, cells were treated with DGAT inhibitors or agonists (PPARα agonist, GW7647, Cayman Chemical Company, 10008613; Forskolin, Millipore-Sigma, F3917; LXR agonist, T0901317, Millipore-Sigma, T2320) for 16 h. The reporter plasmid containing a cAMP response element (CRE; Promega, E8471) was used as a control. PPARs expression vectors were co-transfected with the reporter and control plasmid for 36 h. Cells were then washed with PBS, lysed, and subjected to luciferase activity measurement. Luciferase activities were measured using the Dual-Luciferase Reporter Assay System (Promega, E1910) on a Tecan multimode plate reader. Reporter activity was calculated as the ratio of firefly luciferase to Renilla luciferase to control for transfection efficiency.

### Oxygen consumption measurements

Huh-7 cells were seeded in the XF96 cell-culture microplate (Agilent Technologies) in RPMI supplemented with 10% FBS and settle the plate in the culture hood for 45 min. The following day, cells were treated with DGAT inhibitors for 16 h in RPMI supplemented with 5% lipid-depleted serum. Prior to the assay, cells were washed twice with standard assay medium by supplementing Seahorse XF RPMI medium with 10 mM glucose, 1 mM pyruvate, and 2 mM glutamine and then equilibrated for 1 h at 37°C in a non-CO2 incubator. Etomoxir, Oligomycin, FCCP and Rotenone/Antimycin A stocks were prepared according to the manufacturer’s instructions. OCRs were measured using a Seahorse XFe96 Extracellular Flux Analyzer (Agilent Technologies). The OCR was recorded six times after etomoxir (Cayman Chemical Company, 11969) injection. After completion of the assay, protein content was quantified using a BCA assay for normalization. For acute response to etomoxir measurements, changes in OCR (ΔOCR) were calculated relative to pre-injection baseline values.

### Generation of SREBP1 KO cells with CRISPR/Cas9-mediated gene deletion

SREBP1 KO Huh-7 cells were generated using CRISPR/Cas9 gene editing method^81^. A guide RNA (gRNA) targeting exon 2 of the *SREBP1* locus (5′-CAGCATAGGGTGGGTCAAAT-3′) was designed and cloned into pSpCa9(BB)-2A-Puro (PX459) V2.0 (Addgene 62988) to direct Cas9-mediated cleavage. After transfection, cells were selected with 2 μg/ml puromycin (Thermo Fisher Scientific, A1113803) for 48 h. Puromycin-resistant cells were then trypsinized, serially diluted, and seeded into 96-well plates to isolate single-cell-derived colonies. Positive KO clones were validated by western blot analysis using an anti-SREBP1 antibody and by genomic DNA sequencing (sense: 5′- ACGCACCCTCTTCTCCTT-3′, antisense: 5′- TGTTCCCGGGAGGGCTT -3′).

### Chromatin immunoprecipitation (ChIP)-seq and analysis

Cells were cross-linked with 1% formaldehyde (Thermo Fisher Scientific, 28906) for 10 min at room temperature, and the reaction was quenched by 0.125 M glycine. Fixed cells were washed twice with PBS, collected and lysed in SDS buffer. Nuclei were resuspended in immunoprecipitation buffer and sonicated (Covaris E220) to obtain 200–300-bp DNA fragments. Immunoprecipitation reactions were pre-cleared with 50 μL of Protein G Dynabeads (Thermo Fisher Scientific, 10004D) for 2 h at 4°C and then incubated overnight at 4°C with 5 μg of SREBP1 antibody (Cell Signaling Technology, 95879), H3K4me3 antibody (Epicypher, 13-0041). For normalization, 5 μL of *Drosophila* spike-in chromatin (Active Motif, 53083) and 2 μL of spike-in antibody (Active Motif, 61686) were added to each reaction. BSA-blocked Protein G Dynabeads were added the next day and incubated for 2 h at 4°C. The samples were washed and then reverse-crosslinked overnight in the elution buffer (1% SDS, 0.1 M NaHCO3) and purified using the ChIP DNA Clean & Concentrator kit (Zymo Research, D5205), following the manufacturer’s instructions. After quantification, ChIP DNA was processed at the MSKCC Integrated Genomics Operation (IGO) core, where libraries were prepared using the KAPA EvoPrep kit (Roche) and sequenced on an Illumina NovaSeq 6000.

Raw sequencing reads were trimmed and filtered for quality and library adapters using TrimGalore, and running Cutadapt and FastQC for read evaluation. Reads were aligned to human assembly hg38 using bowtie2 and deduplicated with Picard MarkDuplicates. A peak atlas was generated using BEDTools, and read counts were obtained with featureCounts. Differential enrichment was calculated using DESeq2. Peak densities were normalized using DESeq2 size factors, and peaks were assigned to genes based on intragenic overlap or nearest transcription start site.

### *De novo* lipogenesis measurements

Huh-7 cells were seeded in 150-mm dishes and treated with DGATi inhibitor the following day. After treatment, the medium was replaced with pre-warmed deuterium-labeled medium (RPMI containing 40% D₂O) (Deuterium oxide, Millipore-Sigma, 151882), and cells were incubated for 6 h. Cells were washed with PBS at room temperature, lysed in 0.3 M KOH in MeOH:IPA:H₂O (45:45:10), collected by scraping into glass vials, and saponified at 65°C for 1 h. Samples were cooled to room temperature, acidified with formic acid, and extracted twice with hexane. The combined organic phases were dried in a SpeedVac (45°C, 120 min) and reconstituted in acetonitrile:MeOH (1:1), and 50 μL of supernatant was transferred for LC-MS analysis.

Chromatographic separation was achieved using Waters Acquity UPLC BEH C18 Column (130Å, 1.7 um, 2.1 x 100 mm) with a guard column (Acquity UPLC BEH C18 Column (130Å, 1.7 µm, 2.1 mm X 30 mm). The mobile phase A was water:acetonitrile (95:5) containing 10 mM ammonium acetate and 10 mM ammonium hydroxide, and the mobile phase B was pure methanol. The elution linear gradient was 10% B at 0 min, 60% at 3 min, and 100% at 10.5 min, and held isocratically until 13 min, 10% at 13.01 min, held until 15 min. The flow rate was 0.3 mL/min. The autosampler was at 4°C. The injection volume was 5 μL. Needle wash was applied between samples using methanol: acetonitrile:water at 40:40:20. The MS used was the Orbitrap Exploris 480 (Thermo Fisher Scientific, San Jose, CA) and scanned from 70 to 1000 m/z with switching polarity. The resolution was 480,000. Fatty acid labeling was analyzed by EI-MAVEN.

### MG132 chase assay

Huh-7 cells grown in 100-mm diameter dishes were pre-incubated with DMSO or DGAT inhibitors for 30 min in RPMI supplemented with 5% lipid depleted serum. Cells were washed once with PBS and chased for 0, 1, 2, 3, or 4 h with or without DGAT inhibitors or the proteasome inhibitor MG132 (Millipore-Sigma,474791), as indicated. After completion of the chase at the indicated time points, the medium was removed, and cells were harvested for western blot analysis.

### Nuclear and membrane fractionation of cells

Cells were harvested by scraping and centrifuged at 1000 × *g* for 5 min at 4°C. Nuclear and membrane fraction were extracted as described^40^ with minor modifications. The resulting cell pellets were resuspended in buffer A (250 mM sucrose, 10 mM Hepes-KOH at pH 7.6, 10 mM KCl, 1.5 mM MgCl2, 1 mM sodium EDTA, 1 mM sodium EGTA) with protease inhibitors. The cell suspension was passed through a cell homogenizer 13 times with 10-μm clearance and centrifuged at 1000 ×g at 4°C for 7 min. The pellet was resuspended in buffer B (20 mM Hepes-KOH at pH 7.6, 0.42 M NaCl, 2.5% (v/v) glycerol, 1.5 mM MgCl2, 1 mM sodium EDTA, 1 mM sodium EGTA) with protease inhibitors and then rotated at 4°C for 1 h. The suspension was centrifuged at 10^5^ g for 30 min at 4°C in a Beckman TLA 120.2 rotor. The resulting supernatant was collected as the nuclear extract. The supernatant from the 1,000 × g spin was further centrifuged at 10^5^ g for 30 min at 4°C to obtain the membrane fraction. The resulting pellet was dissolved in SDS-lysis buffer (10 mM Tris-HCl at pH 7.6, 0.1 M NaCl, 1% SDS, 1 mM EDTA, 1 mM EGTA) with protease inhibitors.

### siRNA transfection

siRNAs targeting human DGATs (DGAT1, Dharmacon, L-003922-00-0005; DGAT2, Dharmacon, L-009333-00-0005) and a non-targeting control (Dharmacon, D-001810-01-05) were purchased from Horizon Discovery. siRNAs were resuspended in RNase-free 1x siRNA Buffer and incubated with gentle shaking for 30 min at room temperature. siRNA transfection was performed by using Lipofectamine RNAiMAX (Thermo Fisher Scientific, 13778075) according to the manufacturer’s instructions. Briefly, cells were seeded into six well plates and incubated overnight. The following day, RNAiMAX and siRNAs were separately diluted in Opti-MEM (Thermo Fisher Scientific, 51985034) and then combined to form transfection complexes. The complex was added to the cells and incubated for 24–48 h.

### Microsome purification

Huh-7 cells grown in 15-cm dishes were scraped into ice-cold PBS containing protease inhibitors and centrifuged at 700 × g for 5 min at 4°C. Cell pellets were resuspended in 1 mL of homogenization buffer (250 mM sucrose, 10 mM Hepes-K pH 7.2, 10 mM KCl, 1.5 mM MgCl₂, 1 mM EDTA, and protease inhibitors) and mechanically lysed by 13 passes through a pre-cooled homogenizer. The homogenate was sequentially centrifuged at 700 × g for 10 min and 8,000 × g for 20 min. The resulting supernatant was collected and centrifuged at 10^5^ g for 45 min. The pellet was washed once with the homogenization buffer and stored in -80°C for lipid extraction.

### Lipid extraction

Cells and tissue homogenates were obtained by snap-freezing/thawing repeatedly and extracted, according to a methyl tert-butyl ether (MTBE)–based extraction method^82^. The protein concentration was determined using a BCA assay, and an amount corresponding to approximately 100 µg of total protein was transferred into Pyrex glass tubes with PTFE-lined caps. Lipids were extracted using a total solvent volume of 1200 µL at a 10:3:2.5 (v:v:v) ratio of MTBE:methanol:water. Briefly, 232 µL of methanol was added to each sample, followed by the addition of internal standards (SPLASH® LipidoMix, Avanti Polar Lipids) prior to extraction. Subsequently, 774 µL of MTBE were added, and samples were vortexed vigorously to ensure homogeneous mixing. Samples were incubated at 4°C with intermittent mixing to maximize lipid extraction. Phase separation was induced by adding 194 µL of Milli-Q water, followed by vigorous vertexing.

Samples were centrifuged at 4°C until clear separation was achieved. The upper organic phase, containing extracted lipids, was carefully collected and aliquots of the organic extract were transferred as follows: 500 µL for global lipidomic analysis and 300 µL for fatty acid analysis, each into separate glass tubes. The collected lipid fractions were dried under a gentle stream of nitrogen until complete solvent evaporation and were then reconstituted in isopropanol:acetonitrile:water (60:35:5, v:v:v) and stored at −80°C until MS analysis. The fatty-acid aliquot was dried and stored at −80°C until further processing, according to the downstream fatty-acid analytical workflow.

### LC-MS/MS lipidomic analysis

Lipidomic analyses were performed using ultra-high-performance liquid chromatography coupled to tandem mass spectrometry (UHPLC–MS/MS), adapted from a published method^83^. Chromatographic separation was carried out on a C30 reverse-phase column (Accucore™ C30, 2.1 × 150 mm, 2.6 μm; Thermo Fisher Scientific) maintained at 55°C, using a Vanquish Horizon UHPLC system (Thermo Fisher Scientific) interfaced with an Orbitrap Exploris 240 MS equipped with a heated electrospray ionization source. Each sample (5 μL) was analyzed in both positive and negative ionization modes.

The mobile phase consisted of solvent A, composed of water:acetonitrile (60:40, v:v) with 10 mM ammonium formate and 0.1% formic acid, and solvent B, composed of isopropanol:acetonitrile (90:10, v:v) containing the same additives. Lipids were eluted at a flow rate of 0.26 mL/min using the following gradient: an initial isocratic hold at 30% B from −3 to 0 min, followed by a linear increase to 43% B (0–2 min), 55% B (2–2.1 min), 65% B (2.1–12 min), 85% B (12–18 min), and 100% B (18–20 min). The column was held at 100% B from 20–25 min, then returned to 30% B at 25.1 min, followed by re-equilibration from 25.1–28 min.

MS data were acquired in full-scan MS with data-dependent MS/MS over an m/z range of 250–1700, with internal mass calibration enabled using EASY-IC™. Orbitrap resolution was set to 120,000 for MS1 scans and 30,000 for MS2 scans. The maximum injection time was 50 ms for MS1 and 54 ms for MS2, with the RF lens set to 70%. Source parameters included an ion transfer tube temperature of 300°C, vaporizer temperature of 275°C, sheath gas at 40 units, auxiliary gas at 10 units, and sweep gas at 1 unit. Spray voltages were set to 3250 V in positive mode and 2500 V in negative mode.

Tandem MS spectra were generated using higher-energy collisional dissociation with stepped normalized collision energies of 15%, 25%, and 35%. Data-dependent acquisition was performed with a cycle time of 1.5 s, an isolation window of 1.0 m/z, an intensity threshold of 1.0 × 10⁴, and a dynamic exclusion duration of 2.5 s.

Raw data files were processed using LipidSearch software (version 5.1, ThermoFisher Scientific, OPTON-30879) for lipid identification and alignment, applying a precursor mass tolerance of 5 ppm and a product ion tolerance of 8 ppm. Additional filtering, quality control, and normalization were performed using an in-house data-processing platform (Lipidcruncher^84^). Semi-targeted lipid quantification was achieved by normalizing the area under the curve of individual lipid species to the corresponding internal standards, followed by normalization to the total quantified protein content.

### Targeted LC–MS/MS analysis of free fatty acids

Free fatty acids were analyzed using a targeted UHPLC–MS/MS approach. After MTBE extraction, the organic phase designated for fatty acid analysis was dried under a gentle stream of nitrogen and reconstituted in an LC-MS-compatible solvent consisting of 90% acetonitrile:10% water containing 2 mM ammonium acetate.

Chromatographic separation was performed on a reversed-phase Waters ACQUITY UPLC BEH C18 column (150 × 2.1 mm, 1.7 μm particle size) maintained at 50°C, using a Vanquish UHPLC system (Thermo Fisher Scientific). An isocratic mobile phase composed of 90% acetonitrile and 10% water, supplemented with 2 mM ammonium acetate, was employed throughout the analysis. The flow rate was set to 0.21 mL/min, with a total run time of 15 min. Samples were injected at a volume of 1 μL, and the autosampler temperature was maintained at 10°C.

Mass spectrometric detection was carried out on a high-resolution orbitrap mass spectrometer (Thermo Fisher Scientific) equipped with a heated electrospray ionization source, operated in negative ionization mode. Source parameters included an ion transfer tube temperature of 300°C, vaporizer temperature of 275°C, and a spray voltage of 2500V. Sheath gas, auxiliary gas, and sweep gas were set to 40, 10, and 1 arbitrary units, respectively, with the RF lens set to 70%.

A targeted acquisition method was implemented by defining a pre-specified inclusion list of exact m/z values corresponding to endogenous free fatty acids. Full-scan MS spectra were acquired in the Orbitrap analyzer at a resolution of 120,000 over an m/z range of 100–500, while data-dependent MS/MS acquisition was restricted to the targeted precursor ions specified in the inclusion list. The acquisition cycle time was set to 1.5 s, with an isolation window of 1.0m/z. Fragmentation was performed using higher-energy collisional dissociation with stepped normalized collision energies of 25%, 30%, and 35%, and MS/MS spectra were acquired at a resolution of 30,000. Dynamic exclusion was enabled with a duration of 4 s, and EASY-IC™ was used for internal mass calibration during MS1 acquisition.

Identification of free fatty acids was based on accurate precursor mass, retention time, and characteristic MS/MS fragmentation patterns. Authentic fatty acid standards were analyzed under identical chromatographic and mass spectrometric conditions to establish reference retention times and confirm mass accuracy, and these parameters were used to validate endogenous fatty acid identifications. Mass accuracy was maintained within ≤5 ppm for precursor ions.

Raw LC-MS/MS data were processed using Xcalibur (Thermo Fisher Scientific, OPTON-30965) and LipidSearch, with targeted peak extraction performed using predefined m/z and retention time windows derived from the standards. Quantification was carried out by integrating extracted ion chromatogram (XIC) peak areas for each targeted fatty acid species, followed by normalization to internal standards and sample input, as appropriate.

### ER-laurdan staining and multiphoton microscopy imaging

Huh-7 cells were seeded into 35-mm dishes with 14-mm glass-bottom (MatTek, P35G-1.5-14-C) and cultured in RPMI supplemented with 10% FBS. After the indicated treatments (DGAT inhibitors for 4h or 16h, 100 μM Ethanolamine (Millipore-Sigma, 398136) for 6h and 10 μM SCD1 inhibitor (Cayman Chemical Company, 29421) for 16h), cells were washed once with PBS prior to staining. ER-laurdan (Avanti Polar Lipids, 880197) was dissolved in DMSO to prepare a 5 mM stock solution. Cells were stained with 5 μM ER-laurdan for 20 min in phenol red–free RPMI, then washed once with PBS and three times with phenol red–free RPMI containing 5% lipid-depleted serum.

For generalized polarization (GP) measurements, microscopy experiments were performed using STELLARIS 8 DIVE multiphoton equipped with a HC PL IR APO 40x/1.1 water lens. ER-laurdan was excited with 780-nm laser light, and emission was collected with two 4Tune tunable spectral HyD non-descanned detectors. Fluorescence was detected from two spectrum windows: Ch1 (409–463 nm) for ordered channel detection and Ch2 (473–516 nm) for disordered channel detection^46^. The GP value was calculated at each pixel by 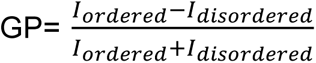 as described^85^. All GP images were corrected using a G-factor determined at the ER-laurdan working concentration used for cell staining. GP value analysis was performed by Amira (Thermo Scientific) and Fiji/ImageJ.

### MINFLUX live cell single-molecule tracking

Huh-7 cells were seeded into 35-mm dishes with 14-mm glass-bottom and transfected with pTK-Halo-truncated CYBT5 and mEmerald-tagged Sec61b for 8 h. Transfected cells were then treated with DGAT inhibitors for 16 h. Janelia Fluor dyes with HaloTag JFX554 and JFX650 were generous gifts from Luke Lavis (Janelia Research Campus). For single-molecule labeling, Huh-7 cells were first labeled with 50 pM JFX650 for 20 min and washed three times with phenol red–free medium (10 min each). Cells were then labeled with 100 nM JFX554 for 1 h and washed three times prior to imaging.

MINFLUX data were acquired on a commercial Abberior Instruments MINFLUX system (Abberior Instruments GmbH) equipped with a 100x/1.4 NA magnification oil immersion lens. A 640-nm continuous-wave laser was used to excite JFX650-labeled CYTB5 in both confocal and MINFLUX mode, while a 488-nm pulsed laser was used for imaging ER marker (Sec61) and a 550 nm laser for imaging the remaining CYTB5 population in confocal mode. Images were acquired with a pixel size of 70 nm. Single-molecule tracking was performed for 30 s using the standard 2D tracking sequence provide by the Abberior Instruments Inspector software. Confocal images of the ER marker were acquired before and after MINFLUX tracking as a spatial reference.

### MINFLUX data analysis and apparent diffusion calculation

MINFLUX single-molecule localization data were acquired across multiple experimental sessions and processed using custom Python scripts to extract track coordinates and assign localizations relative to LD structures. Raw .npy files containing X, Y, and time values were imported into Pandas, and spatial coordinates were converted to µm using instrument-specific pixel scaling (70 nm/pixel) and offset parameters obtained from metadata or auxiliary parameter files. Tracks with fewer than 500 localizations, limited spatial dispersion, or minimal net displacement were excluded to ensure trajectory quality. Processed trajectories were visualized by overlaying tracks onto ER confocal images, and tracks from all biological replicates were combined into a final dataset for downstream analysis.

Localization precision was estimated from the standard deviation of frame-to-frame positional changes in the X and Y axes. To assess local diffusivity along individual trajectories, a sliding-window mean squared displacement (MSD) analysis was performed using a window corresponding to ∼5 ms, determined from the median inter-frame interval. Within each window, squared displacements were calculated across incremental time lags relative to the initial localization, and MSD curves were generated. Apparent diffusion coefficients (D) were estimated from the MSD slope using the Brownian diffusion relation D=MSD/(4t). Mean D values from all windows within a track were aggregated to generate per-track diffusion measurements for downstream analysis.

### Statistics and reproducibility

Statistical analyses were performed using GraphPad Prism 10. Data were presented as mean ± standard deviation, unless otherwise indicated. Comparisons between two groups were performed using two-tailed unpaired Student’s *t*-tests. For two-variable datasets, we used a two-way ANOVA analysis in GraphPad Prism. The number of biological replicates for *in vivo* and *in vitro* experiments are indicated in the figure legends. Statistical differences are indicated by the p-values mentioned in the figures.

---

**Extended Data Figure 1.**
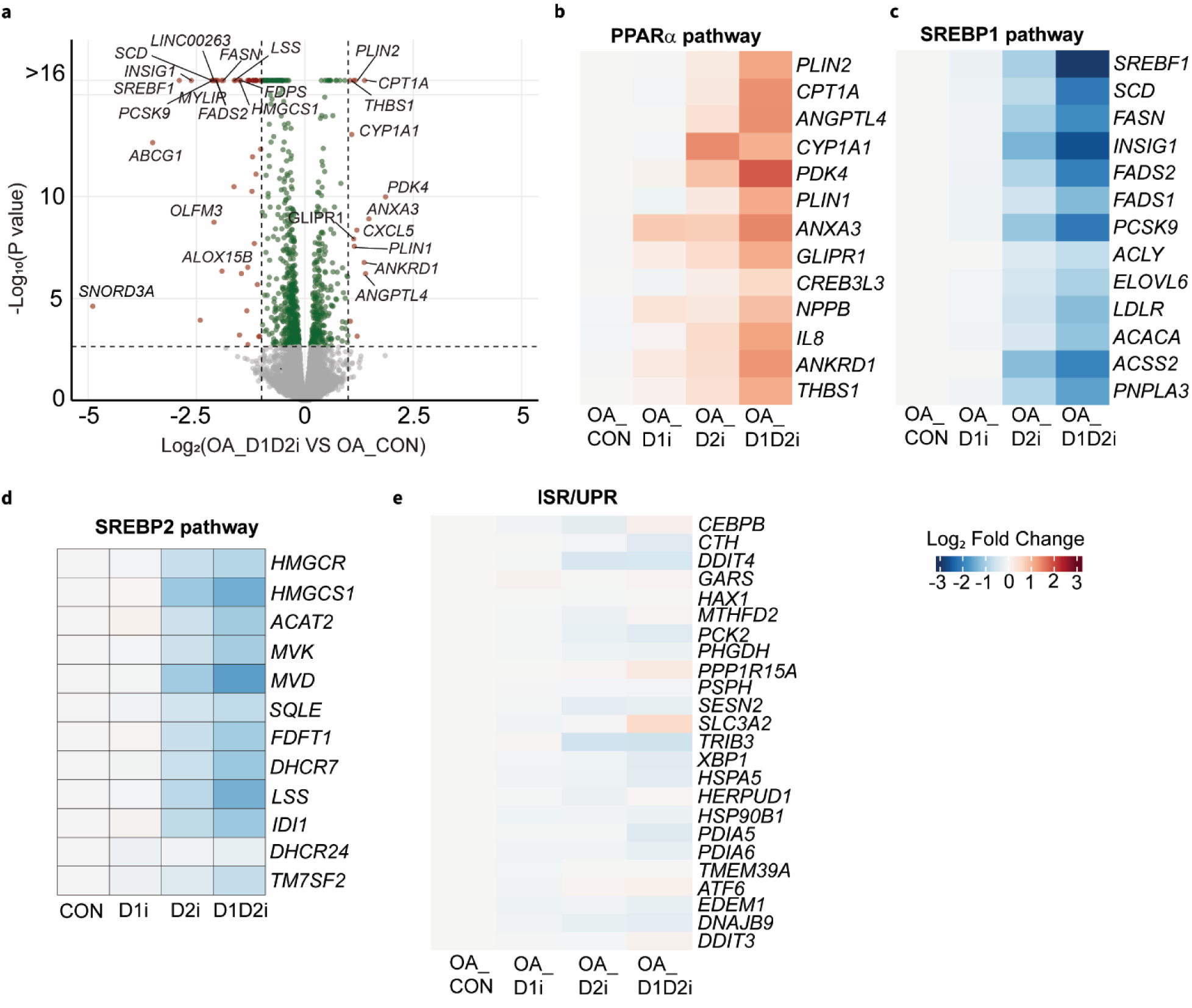
Inhibiting TG synthesis under oleic acid supplement triggers transcriptional responses in SREBP1 and PPARα pathways. **(a)** Volcano plot showing differentially expressed genes between combined DGAT1 and DGAT2 inhibition (OA_D1D2i) and DMSO (OA_CON) conditions in the presence of 200 µM oleic acid (OA). Genes with adjusted p values < 0.05 and absolute fold-change > 1 were considered significantly differentially expressed. Significantly altered genes are shown in red. n = 3 for each treatment. **(b)** Heatmap showing relative gene expression levels of genes involved in the PPARα pathway derived from RNA-seq analysis. **(c)** Heatmap showing relative gene expression levels of genes involved in the SREBP1 pathway derived from RNA-seq analysis. Huh-7 cells were treated with DMSO (OA_CON), DGAT1 inhibitor (OA_D1i, 10 µM), DGAT2 inhibitor (OA_D2i, 10 µM), or combined DGAT1 and DGAT2 inhibition (OA_D1D2i, 10 µM each) in the presence of 200 µM oleic acid for 16 h. **(d)** Heatmap showing relative gene expression levels of genes involved in the SREBP2 pathway. **(e)** Heatmap showing relative gene expression levels of integrated stress response (ISR) and unfolded protein response (UPR) marker genes following 16 h of DGAT inhibitors treatments in the presence of OA.

**Extended Data Figure 2.**
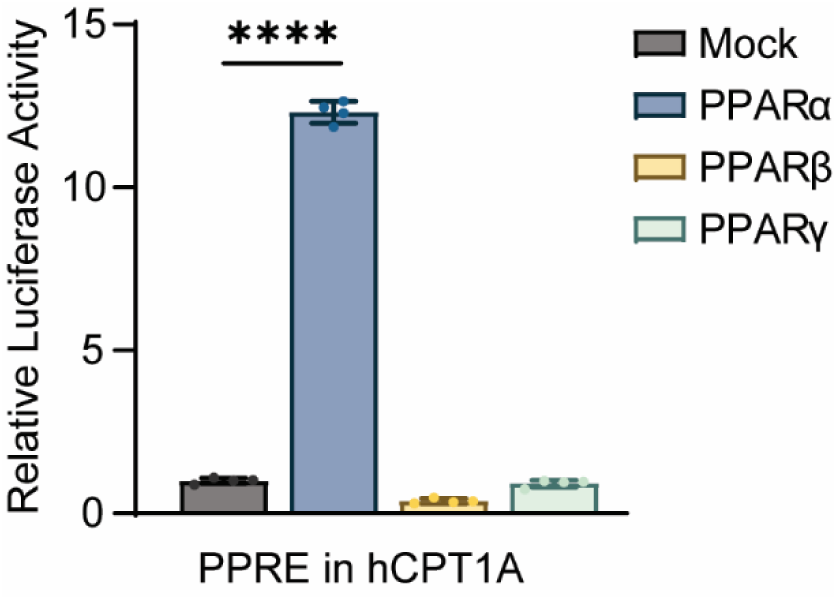
Effects of three PPAR isoforms (α, β and γ) on the transcriptional activity of PPRE in human CPT1A. Huh-7 cells were transfected with the reporter of PPRE in human CPT1A and PPARs expression plasmid for 36 h. Cells were washed and lysate for dual-luciferase reporter assay. Data are presented as mean ± SD, n = 4 for each treatment. Statistical significance was evaluated by unpaired two-tailed student’s t test. ∗∗∗∗p < 0.0001.

**Extended Data Figure 3.**
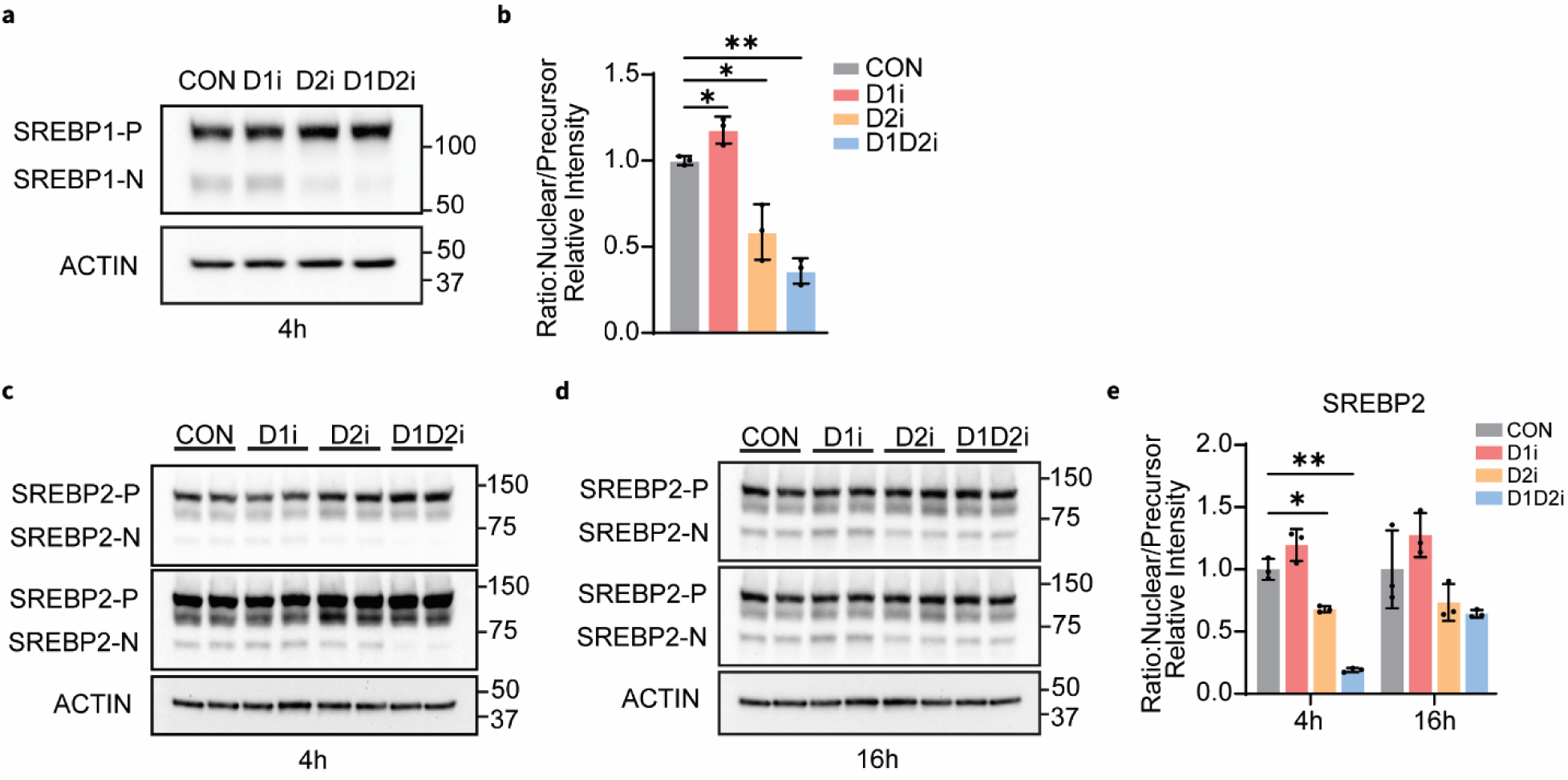
Impaired TG synthesis selectively disrupts SREBP1 processing. **(a)** Western blot analysis of SREBP1 in whole-cell lysates from Huh-7 cells treated with DMSO (CON), DGAT1i inhibitor (D1i), DGAT2 inhibitor (D2i), or combined DGAT1 and DGAT2 inhibition (D1D2i) for 4 h. Actin was used as loading control. **(b)** Bar graphs showing quantification of the relative ratio of nuclear (N) to precursor (P) SREBP1. Mean ± SD, n = 3, ∗p < 0.05; ∗∗p < 0.01. Statistical significance was evaluated by unpaired two-tailed student’s t test. **(c-d)** Western blot of SREBP2 in whole-cell lysates from Huh-7 cells treated with DMSO (CON), DGAT1i inhibitor (D1i), DGAT2 inhibitor (D2i), or combined DGAT1 and DGAT2 inhibition (D1D2i) for 4 h (**c**) and 16 h (**d**). **(e)** Bar graphs showing the relative nuclear-to-precursor (N/P) ratio of SREBP2. Mean ± SD, n = 3, ∗p < 0.05; ∗∗p < 0.01.

**Extended Data Figure 4.**
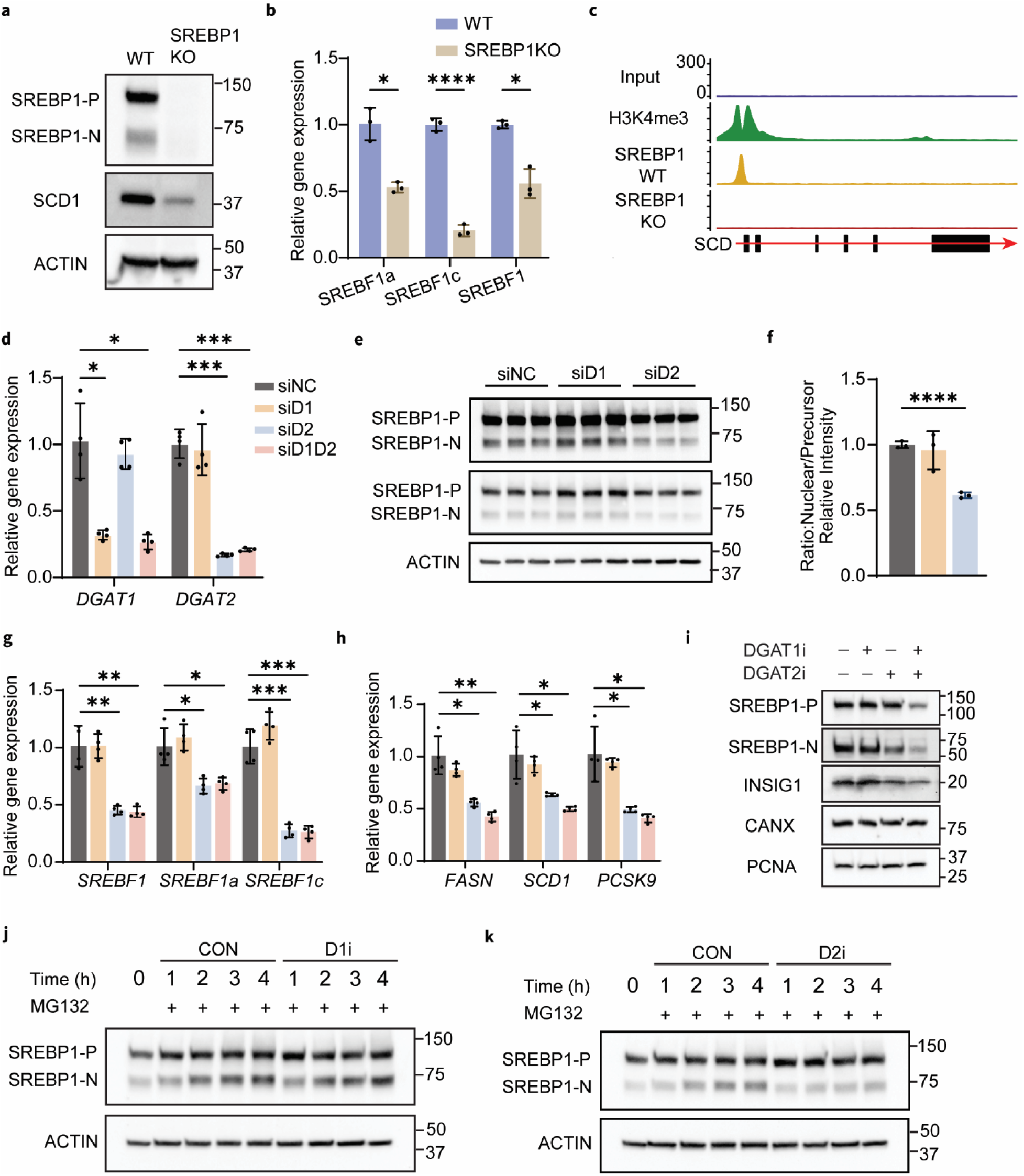
Impaired TG synthesis consistently suppresses SREBP1 processing and expression in Huh-7 cells. **(a)** Validation of SREBP1 KO in Huh-7 cells by western blot analysis of SREBP1 and SCD1. Actin was used as loading control. **(b)** Relative expression of *SREBF1*, *SREBF1a*, and *SREBF1c* in WT and SREBP1 KO Huh-7 cells. Mean ± SD, n = 3. ∗p < 0.05; ∗∗∗∗p < 0.0001. Statistical significance was evaluated by unpaired two-tailed student’s t test. **(c)** Representative ChIP-seq tracks at the SCD locus. Shown are input control, H3K4me3, and SREBP1 ChIP-seq signals in WT and SREBP1 KO Huh-7 cells. **(d)** Relative gene expression of *DGAT1* and *DGAT2* in Huh-7 cells transfected with DGAT-targeting siRNA for 48 h. Mean ± SD, n = 4. ∗p < 0.05; ∗∗∗p < 0.001. **(e-f)** Western blot analysis (**e**) and quantification (**f**) of SREBP1 in whole-cell lysates of Huh-7 cells transfected with DGAT-targeting siRNA for 48 h. Mean ± SD, n = 3. ∗∗∗∗p < 0.0001. **(g-h)** Bar graph showing relative gene expression of *SREBF1* and its target genes in Huh-7 cells transfected with DGAT-targeting siRNA. Mean ± SD, n = 4. ∗p < 0.05; ∗∗p < 0.01; ∗∗∗p < 0.001. **(i)** Western blot analysis of SREBP1 precursor (SREBP1-P) and INSIG1 in membrane fractions and nuclear SREBP1 (SREBP1-N) in nuclear fractions. Calnexin and PCNA were used as membrane and nuclear markers, respectively. **(j-k)** Western blot analysis of SREBP1 in whole-cell lysates of Huh-7 cells treated with DMSO (CON) or DGAT inhibitors (DGAT1 inhibitor, D1i; DGAT2 inhibitor, D2i) in the presence of MG132 at the indicated time points.

**Extended Data Figure 5.**
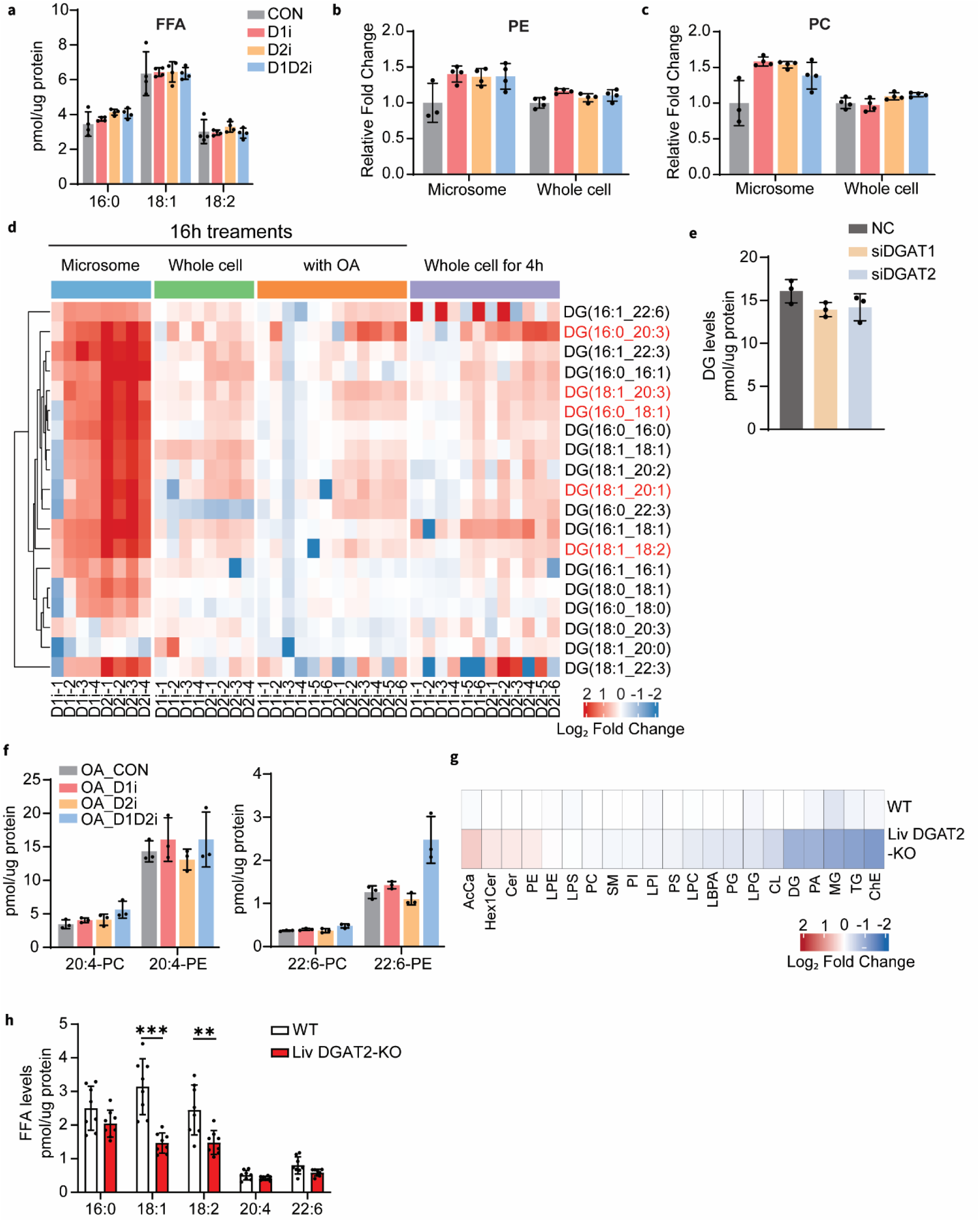
Impaired TG synthesis changes lipid profiles in Huh-7 cells and murine liver. **(a)** Bar graph showing free fatty acids levels (C16:0, C18:1 and C18:2) in whole-cell lysates of Huh-7 cells treated with DGAT inhibitors for 16 h. Data are presented as mean ± SD. **(b-c)** Bar graphs showing relative fold-changes in phosphatidylethanolamine (PE) and phosphatidylcholine (PC) levels in microsome and whole-cell lysates of Huh-7 cells treated with DGAT inhibitors for 16 h. Data are presented as mean ± SD. **(d)** Heatmap analysis of fold-change of diacylglyceride (DG) levels in Huh-7 cells across multiple treatment conditions, normalized to the DMSO (CON) control group. DG species showing consistent changes are highlighted in red. **(e)** Bar graph showing total DG levels in whole-cell lysate of Huh-7 cells transfected with DGAT-targeting siRNA. Mean ± SD, n = 3. **(f)** Bar graphs showing the abundances of phospholipids containing arachidonic acid (20:4) or docosahexaenoic acid (22:6) in whole-cell lysates of Huh-7 cells treated with DGAT inhibitors in the presence of 200 μM oleic acid for 16 h. **(g)** Heatmap analysis of fold-change of major lipid class abundances in liver tissues of WT and liver-specific DGAT2 knock out (DGAT2-KO) mice, normalized to control. Data plotted as mean of the log2 (fold-change). Mean ± SD, n = 8. **(h)** Free fatty acid levels in in liver tissues of WT and liver-specific DGAT2-KO. Mean ± SD, n = 8. ∗∗p < 0.01; ∗∗∗p < 0.001.

**Extended Data Figure 6.**
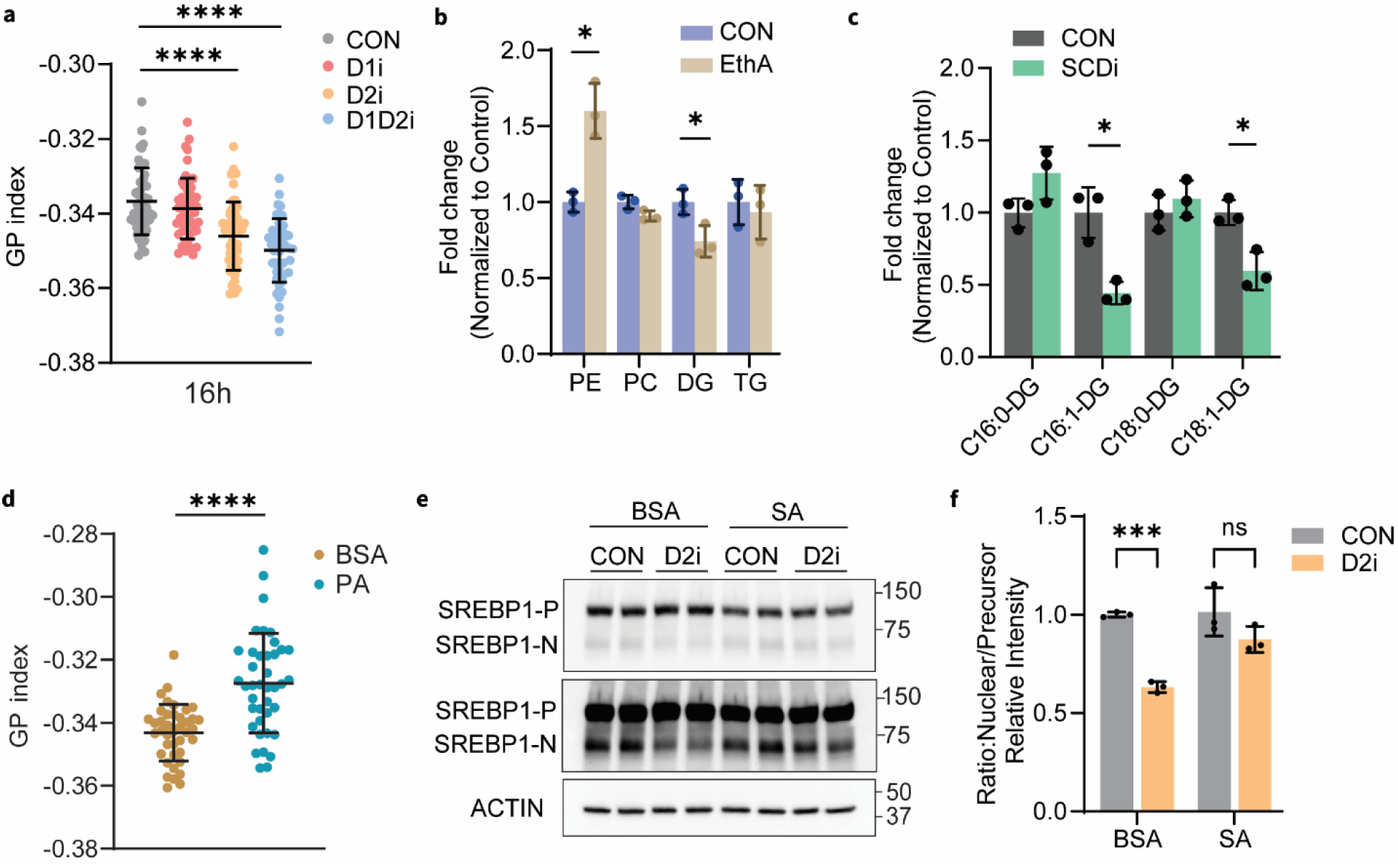
SREBP1 processing is regulated by ER membrane fluidity. **(a)** Quantification of GP index of Huh-7 treated with DMSO or DGAT inhibitors for 16 h. Mean ± SD, n = 50 cells, ∗∗∗∗p < 0.0001. Statistical significance was evaluated by unpaired two-tailed t-test. **(b)** Bar graphs showing the relative fold-change of phospholipids (PC and PE) and neutral lipids (DG and TG) in the Huh-7 cells incubated with ethanolamine (EthA, 100 μM) for 6 h. Mean ± SD, n = 3. ∗p < 0.05. Statistical significance was evaluated by unpaired two-tailed t-test. **(c)** Bar graphs showing the relative fold-change of DG containing saturated fatty acids (C16:0 or C18:0) or monounsaturated fatty acids (C16:1 or C18:1) in the Huh-7 cells incubated with SCD1 inhibitor for 16 h. Mean ± SD, n = 3. ∗p < 0.05. (d) Quantification of GP index of Huh-7 incubated with BSA or palmitate (PA, 200 μM) for 4 h. Mean ± SD, n = 40 cells, ∗∗∗∗p < 0.0001. **(e-f)** Western blot analysis and quantification of SREBP1 in whole-cell lysates. Huh-7 cells were pretreated with BSA or stearic acid (SA, 200 μM) for 4 h and then treated with DMSO or DGAT2 inhibitor in the presence BSA or SA. Mean ± SD, n = 3. Statistical significance was determined by two-way ANOVA, followed by Šídák’s multiple comparisons test for selected pairwise comparisons, ∗∗∗p < 0.001.

## Notes

### Competing Interest Statement

The authors have declared no competing interest.

### Summary of Updates

This version of the manuscript has been revised to update the following (Supplemental files updated).

